# Cognitive sequences in obsessive-compulsive disorder are supported by frontal cortex ramping activity

**DOI:** 10.1101/2024.07.28.605508

**Authors:** Hannah Doyle, Nicole C.R. McLaughlin, Sarah L. Garnaat, Theresa M. Desrochers

## Abstract

Completing sequences is a routine part of daily life. Many are abstract, defined by a rule governing the order rather than the identity of individual steps (e.g., getting dressed). In obsessive-compulsive disorder (OCD), excessive ritualistic behaviors suggest a disruption in abstract sequence completion. Executing abstract sequences requires at least two levels in a hierarchy of cognitive control: abstract sequential control (tracking steps) and task switching (shifting between tasks). While task switching has been studied in OCD, little is known in a sequential context. Understanding both hierarchical control types is key to uncovering how abstract sequences with nested task switches are processed in OCD. Previous studies showed that the rostrolateral prefrontal cortex (RLPFC) supports abstract sequence monitoring in healthy individuals with an increase in activity across each sequence, a dynamic known as “ramping”. Ramping outside the RLPFC is potentially indicative of other sequence-related processes such as progress towards a goal and increasing working memory load. Therefore, we hypothesized that abstract sequential control deficits would correspond to altered ramping dynamics in RLPFC and other cortical regions. Second, we predicted task switching deficits in OCD, coupled with altered activity in cortical regions canonically implicated in task level control. We found partial support for both hypotheses. Abstract sequential control did not show behavioral differences in OCD but did show increased overall ramping in the anterior cingulate cortex (ACC) and superior frontal sulcus (SFS) and ramping differences in additional, novel cortical regions according to abstract sequence complexity. In contrast, behavioral differences were observed for task switching in OCD without neural differences between the groups. Together, these results suggest a group of areas support sequential control differentially in OCD than in healthy controls, despite behavioral similarity, and that this observation is likely not the result of neural deficits in task switching. These findings thus provide insight into OCD during complex behaviors more similar to daily life where sequence and task level control are intertwined and may inform future potential treatment.

## Introduction

A relatively prevalent psychiatric disorder, obsessive-compulsive disorder (OCD), is defined by repetitive obsessions (persistent and intrusive thoughts, feelings, or images) and associated compulsions (Veale & Roberts, 2014). Behaviorally, compulsions present as repetitive and unnecessary, often ritualistic, actions or thoughts. Common examples of compulsions include repeatedly checking items or cleaning rituals. Although they can differ between individuals, compulsions in OCD can be categorized broadly as sequences of repeated actions or thoughts that are illogical or excessive to the situation. Therefore, it is important to investigate the neural contributions to the sequential nature of OCD behavior to understand the basis of its pathology.

Compulsive behaviors in OCD, such as counting in groupings of a certain number (e.g., groups of five) (Menon, 2013), or dressing and re-redressing in the morning (Uvais & Sreeraj, 2016), suggest a potential for dysfunction in the ability to track task sequences. In daily life, sequences define the way humans organize their lives, often establishing a scaffold we can use to help achieve our goals. Many such sequences can be considered abstract, in that they are defined by a rule governing a series of operations over a series of tasks rather than by the identity of the operations themselves (Desrochers et al., 2022). For example, the abstract sequence of cooking pasta may be guided by the structure of a recipe (e.g., boil water for noodles, chop the vegetables, grate the cheese), with the flexibility of using tomatoes from the garden or the store without disrupting the process. It is possible that dysfunctional neural circuitry related to abstract sequence behavior underlies OCD pathology, since many compulsions manifest as repeated or ritualistic abstract sequences.

Abstract task sequences consist of a hierarchy of at least two levels of cognitive control required for their completion. The top level of the hierarchy is abstract sequential control, which is the cognitive flexibility and maintenance needed to keep track of each step with an end goal in mind (Desrochers et al., 2022; Lashley, 1951). The second level of the hierarchy is task or sub-goal control, a more general type of control required to cease one task and switch to the other, often referred to as task switching (Monsell, 2003). Abstract sequences therefore require the exertion of cognitive control over tracking progress made towards a goal and over individual task transitions.

Both abstract sequential control and task switching have established behavioral markers. In abstract sequencing, healthy participants exhibit significantly increased reaction times at sequence onset compared to later sequence positions (Desrochers et al., 2015, 2019; Schneider & Logan, 2006). This behavioral effect is known as the initiation cost and is thought to measure control at the abstract sequential hierarchy level. The symptomatology of OCD suggests initiation costs may be increased as a result of impaired abstract sequential control in these patients, though there have been few direct studies. The behavioral marker for task level control has also been established. During task switching, significantly higher reaction times occur at task switches compared to repeats in healthy participants (Monsell, 2003). Evidence of task level control deficits in OCD is mixed with some studies reporting switch cost deficits (Gu et al., 2008; Remijnse et al., 2013), and another concluding there were no task level deficits using a similar task (Moritz et al., 2004). In paradigms that combine hierarchical levels of control, evidence is also mixed as to the effects of the combination. In healthy participants, predictable task sequences can improve task switching reaction times (Schneider & Logan, 2006), while conversely, conflicting higher order rules can increase reaction times on embedded switches (Mayr & Bryck, 2005). In OCD, one study found that in a dual-task paradigm that combined inhibitory control with switching tasks, OCD participants had more interference on switches, resulting in higher switch costs (Demeter et al., 2017). These studies underscore the importance of examining markers of different levels of behavioral control together to better understand OCD symptoms as they relate to abstract and task levels of sequential control.

Neural markers at the level of abstract sequential control are established through two distinct dynamics, onset and ramping activity. Ramping refers to a monotonic increase in activity across a sequence. In healthy participants, ramping occurs in the rostrolateral prefrontal cortex (RLPFC), a region necessary for successful sequence task performance, reflecting the extended monitoring or tracking of progress through abstract sequences (Desrochers et al., 2015, 2019; McKim & Desrochers, 2022). By compliment, onset activity, the amplitude of the response at the start of each stimulus, also occurs in the RLPFC and can index how this region contributes to sequential control that is more trial specific and not extended across entire sequences. In OCD, reduced onset activity occurs in prefrontal regions, such as the ventromedial (Gu et al., 2008) and dorsolateral (Gu et al., 2008; Remijnse et al., 2013; van den Heuvel et al., 2005) cortex. Given the role of these regions in general executive functioning, it is plausible that hypoactivity may occur in the RLFPC during sequential control tasks. If abstract sequential control is disrupted in OCD, we would expect differences in RLPFC ramping, and potentially in onset activity as well. Ramping and onset dynamics particularly in the RLPFC, are therefore important to index abstract sequential control.

Beyond the RLPFC, ramping has been observed in a network of areas during sequential tasks (Desrochers et al., 2015, 2019; McKim & Desrochers, 2022). Because this dynamic emerges in these areas in the context of a sequential task, these processes are also likely sequence related. For instance, ramping in the ACC may index growing demands for performance monitoring or error detection (Blanchard et al., 2015), or reflect the tracking of motor responses within a sequence in the supplementary motor area (SMA) (Bonini et al., 2014) or relate to accumulating working memory load in temporal lobe regions (Lundqvist et al., 2018). By assessing ramping throughout the task, we aim to dissociate RLPFC-specific signals of abstract sequence progression from ramping that potentially reflects complementary processes in other cortical regions.

At the task level of control, differences in onset dynamics could indicate altered task switching in OCD. This dynamic is the most direct way to measure task switching effects in the sequential task since it is sensitive to trial-by-trial differences. Although evidence is mixed, several studies show hypoactivity in control regions like the ventromedial PFC (Gu et al., 2008) and the DLPFC during task switching (Gu et al., 2008; Remijnse et al., 2013; van den Heuvel et al., 2005) in OCD participants. Task switching engages a broad network of cortical regions, so we may expect to see hypoactivity in such regions in OCD in the present study related to the task level of control. Together both ramping and onset dynamics can be used across the two hierarchical levels of control to gain insight into the neural bases of abstract sequence performance in OCD.

In addition to investigating neural dynamics alone, examining the relation to symptom severity based on categorical diagnoses provides a more comprehensive view of how hierarchical control levels are altered in OCD during abstract sequence processing. Related to general cognitive control at the task level, OCD symptom severity has been found to correspond to delayed reaction times and attentional deficits in one task switching study (Okutucu et al., 2023). Another study similarly reported correlations between OCD symptom severity and task switching error rates (Remijnse et al., 2013). These highlighted studies suggest OCD symptoms may also correlate with prefrontal cortical neural dynamics during abstract sequential control, another part of the broader cognitive control spectrum.

We investigated the neural underpinnings of abstract and task levels of sequential control in participants with OCD and healthy controls (HCs) using functional magnetic resonance imaging (fMRI). We used a behavioral paradigm that assessed both abstract sequence and task level control and examined ramping and onset dynamics. First, at the level of abstract sequential control, we predicted that OCD participants would exhibit deficits reflected in hypoactivity and decreased ramping dynamics in the RLPFC during the task, and potentially in additional cortical regions that support auxiliary cognitive resources utilized in the task. We additionally expected these RLPFC differences to correlate with increased OCD symptom severity. Second, at the level of task control we predicted that OCD participants would exhibit behavioral impairments and that these would be associated with hypoactivity in cortical regions beyond the RLPFC that are typically engaged in task-switching processes. We observed partial support for both hypotheses. At the level of abstract sequential control, we did not observe behavior deficits in OCD, and we did not observe RLPFC activity or ramping differences between groups. However, in contrasts that probe this level, we observed increased ramping in OCD in the pregenual anterior cingulate cortex (rACC) and superior frontal sulcus (SFS) and additional novel cortical regions. At the task control level, OCD participants exhibited differences in error rates but no neural activity differences directly underlying this behavior. These findings overall implicate novel cortical regions for supporting the level of abstract sequential control in OCD and inform current neurobiological models of and future treatments for patients.

## Methods

### Participants

Participants were recruited via online advertising, fliers, and word of mouth. All participants gave informed, written consent and study procedures were approved by the Butler Hospital Institutional Review Board. Initially, 76 participants were recruited to complete the clinical interview and abstract sequencing task. Of these, 16 dropped out or were screened out of the study and therefore did not advance to the fMRI scan, resulting in 60 scanned in total. Two participants were excluded for excessive motion, one due to user error in handling the button box, seven due to poor behavioral performance (overall error rates > 20% on average across all runs and trials). As previous studies using this task have demonstrated, error rates below 20% on average across runs ensured participants were completing the task as instructed (Desrochers et al., 2015, 2019; Schneider & Logan, 2006; Trach et al., 2021). After excluding participants, our final sample size was 50 in total, 25 in the HC group (mean 28.9 yrs (+/-10.7 [SD]); 11 m [14 f]), and 25 in the OCD group (mean 25.8 yrs (+/-8.5 [SD]); 3 m [22 f]). The target recruitment age range for this study was 18 – 55, so we note that the average age in both groups may be lower due to the population in and around the area being skewed towards college-age participants. However, mean ages were not significantly different between groups (independent samples t-test: t(48) = 1.15, p = 0.25), suggesting that the proportion of college-age participants is not different between the groups. Efforts were also made to recruit similarly for each group. More comprehensive demographic information for each participant group is reported in **Table 1**. The original target sample size was 26 in each group based on a power analysis used to determine sample size in a previous study using this paradigm in healthy controls (Desrochers et al., 2015), however, a post-hoc power analysis determined we achieved 78% power given a sample size of 25 in each group for an effect size of 0.5 (Cohen’s d).

**Table 1.** Demographic information for both HC and OCD participant groups. Mean age (in years) and scores for the Y-BOCS and QIDS-SR were reported with +/-1 SD (standard deviation). Note that no Y-BOCS scores were collected for HCs and thus were not reported. Total number of males and females were reported. Percentage identifying of a particular race was reported per group. Note that the total percentage adds to over 100% as participants were allowed to identify as more than one race. The total number identifying as Not Hispanic/Hispanic or Unknown identity were reported.

|  | HC | OCD |
| --- | --- | --- |
| **Age** | 28.9 yrs. (+/- 10.7) | 25.8 yrs. (+/- 8.5) |
| **Sex** | 11 m (14 f) | 3 m (22 f) |
| **Race** | 87% White, 13% Asian, 1% Black | 84% White, <1% Asian, 12% Black |
| **Ethnicity** | 17(6) Not Hispanic (Hispanic) | 20(4)(1) Not Hispanic (Hispanic)(Unknown) |
| **Y-BOCS** | N/A | 22.28 (+/- 4.09) |
| **QIDS (SR)** | 4.39 (+/- 3.04) | 9.00 (+/- 4.44) |

Note for the HC group: Age and Sex information are reported for all 25 in the HC group (total number in this group). In the HC group, 2/25 participants completed this fMRI study prior to the clinical interview installment within the protocol, so additional demographic data (Race, Ethnicity, and clinical scores) were not collected and therefore not reported for these two participants.

Inclusion criteria for the healthy control group were as follows: 18 - 55 years of age, right-handed, ability to communicate in English to perform study procedures and provide consent. OCD group inclusion criteria followed that of the healthy control group with the following additions: current DSM-5 diagnosis of OCD and Y-BOCS score of equal to or greater than 16, no use or stable psychiatric medication use for 6 weeks prior to study enrollment, limited to serotonin reuptake inhibitors and PRN use of benzodiazepines. Healthy control group exclusion criteria were as follows: current psychiatric diagnosis, lifetime diagnosis of psychotic disorder, bipolar mood disorder or OCD, active suicidal ideation, significant neurological pathology, use of psychiatric medications, contraindications to MRI scan (e.g., ferromagnetic implants, pregnancy, or other conditions that pose safety risk). OCD exclusion criteria were as follows: active problematic substance use, lifetime diagnosis of psychotic disorders, history of bipolar disorder/mania, clinically significant hoarding symptoms, active suicidal ideation, significant neurological pathology, and contraindication to MRI scans.

Each participant completed an interview and an in-person fMRI scan. The clinical interview visit consisted of completing informed consent, and administration of clinical interviews and self-report measures (as described below). Participants could only proceed to the fMRI portion if they were still eligible for the experiment after this first study visit. The second session was an fMRI scan conducted at the Brown University MRI Research Facility. To overview the scan session, participants were first trained on the task and then completed 5 runs of the task in the MRI scanner. Participants were compensated $25 for the first session and $75 for the second. Participants who were ineligible for the fMRI portion were only compensated for the clinical interview visit.

### Measures

The cognitive task and clinical interviews were administered by trained evaluators (see description of, below), and participants completed additional self-report measures.

#### Clinician Administered

*Structured Clinical Interview for DSM-5 (SCID-5)* (First et al., 2017) is an evaluator-administered semi-structured interview to assess for presence or absence of specific psychiatric disorders. In this study, the following selected modules of the SCID-5 were used: mood disorders, anxiety disorders, OCD and related disorders, and trauma-related disorders. Psychotic disorders and hoarding disorder were screened using the SCID-5 and excluded in the present study.

*Yale-Brown Obsessive-Compulsive Scale (Y-BOCS)*(Goodman, Price, Rasmussen, Mazure, Delgado, et al., 1989; Goodman, Price, Rasmussen, Mazure, Fleischmann, et al., 1989): The Y-BOCS symptom checklist is an evaluator-administered measure used to assess presence or absence of common OCD symptoms. The accompanying Y-BOCS severity scale is an evaluator-administered assessment of OCD symptoms severity measured over the past week. The Y-BOCS is considered the gold-standard measure of OCD symptom severity.

#### Self-report

*Alcohol Use Disorders Identification Test (AUDIT)* (Saunders et al., 1993): The AUDIT is a 10-item self-report questionnaire that assesses alcohol consumption, drinking behaviors, and alcohol-related problems. A score of 8 or above was used as a cut-off for men, while a score of 6 or above was used as exclusion criteria for women (Bergman & Källmén, 2002). Scores range from 0 - 40, with a higher score indicating more alcohol use.

*Drug Use Disorders Identification Test (DUDIT)* (Berman et al., 2016): The DUDIT is an 11-item self-report measure that assesses current drug-related problems or drug abuse. A score of 6 or higher was used as an exclusion criterion for men while a score of 2 or higher screened out women in the current study (Berman et al., 2005). Scores range from 0 - 44, with higher scores indicating more drug use.

*Quick inventory of depressive symptomatology (QIDS-SC)* (Rush et al., 2003): The QIDS-SC is a 16-item self-report measure of depression severity. Scores range from 0 - 27, with higher scores indicating more severe symptoms.

### Task Design and Procedure

#### Overview

The abstract sequence task used in this study was used in a previous study of healthy controls (**Figure 1**) (Desrochers et al., 2015) and was based on previous studies of sequential control (Schneider & Logan, 2006). On each trial, participants were presented with a stimulus of varying size (small [3.5 x 3.5 cm] or large [7 x 7 cm]), shape (circle or square), and color (red or blue), for a total of 8 possible stimuli that appeared equally throughout the task and did not repeat on adjacent trials. After each trial was an intertrial interval, displayed as a white fixation cross centered on a black screen, with jittered timing (0.25 - 8 s). Participants were provided 4 seconds on each trial to make a response. Each trial had response options for the color and shape of the stimulus, mapped onto two response pad buttons, corresponding to the index and middle finger of the right hand. Each response option was one shape and color combination (e.g., index finger button maps onto both ‘blue’ and ‘circle’ and the middle finger maps onto ‘red’ and ‘square’). Participants pressed one button per trial to indicate their response. Response options were always shown on the bottom left and right of the screen. Stimulus-response mappings were kept consistent throughout the experiment but were counterbalanced across participants. The frequency of responses to each stimulus and the response repeats (instances when the same finger was used to respond to two trials in a row) were counterbalanced throughout the task.

**Figure 1.**
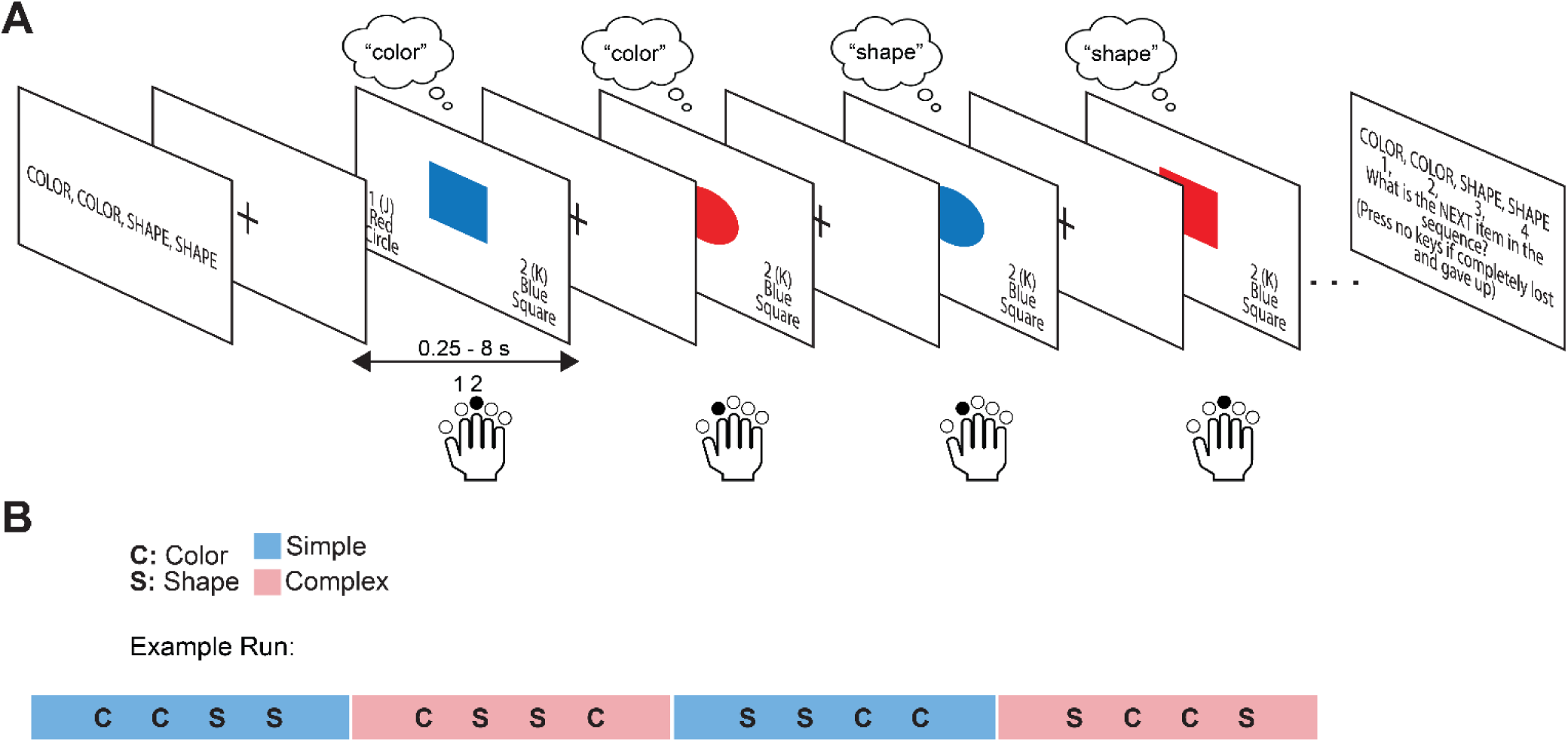
Abstract sequence task schematic. **A.** Example trials in a block for the simple sequence. Each block begins with a screen that instructs the sequence, e.g., “COLOR, COLOR, SHAPE, SHAPE”. Each trial consists of one stimulus presentation where the participant must make the correct categorization decision based on the identity of the stimulus and the position in the sequence. The remembered categorization decision for each item is indicated in a thought bubble and the correct choices for each trial are indicated by black arrows. The stimulus remains on screen until a response is made (max 4 sec). After the response (or response time-out), a fixation cross is displayed for the duration of the intertrial interval (ITI, jittered 25 - 8000 ms). Distance between images is for illustration purposes only and does not represent actual timing. There are 24-27 trials per block, it can end on any position in the sequence, and the block ends with a sequence position question asking, “What is the NEXT item in the sequence?”. **B.** Example run containing four blocks, with each block being a simple (CCSS [color, color, shape, shape]; SSCC [shape, shape, color, color]) or complex (CSSC [color, shape, shape, color]; SCCS [shape, color, color, shape]) sequence. The order of the blocks is counterbalanced across the five runs that each participant performs.

Stimuli were presented in blocks (24-27 trials, so that blocks ended on unpredictable sequence positions, counterbalanced across blocks), and participants completed 4 blocks per run, for 5 runs total. At the beginning of each block, participants were shown a 4-item sequence (5 s), which they used to make a choice on every trial, followed by a fixation screen (1 s). Every block consisted of a sequence that was one of two types: simple (of the pattern AABB; specifically “COLOR COLOR SHAPE SHAPE” or “SHAPE SHAPE COLOR COLOR”) or complex (of the pattern ABBA; specifically “COLOR SHAPE SHAPE COLOR” or “SHAPE COLOR COLOR SHAPE”). Simple sequences contained one embedded task switch (e.g., switching on positions 2 to 3 from “COLOR” to “SHAPE” in the sequence “COLOR COLOR SHAPE SHAPE”) while complex sequences contained two embedded task switches (e.g., switching on positions 1 to 2 from “COLOR” to “SHAPE” in the sequence “COLOR SHAPE SHAPE COLOR”). The number of task switches was equivalent across blocks, so that the probability of occurring switch or repeat trials was equal between blocks of complex and simple sequences. At the end of each block, participants were shown a screen that asked what sequence position they would be on if they were to make a choice on the next trial. Participants responded to this question using one of four buttons on the response pad (excluding the thumb button). The order of simple and complex sequence blocks were counterbalanced across runs.

Participants were trained on an Alienware M17xR4 laptop (Windows 10) using a five-button response pad on four shortened task blocks prior to scanning. Participants completed practice on response pad buttons and then were guided by the experimenter on each trial for the first practice block. Participants performed the remaining practice blocks independently. Performance competency was established by error rates less than 20% overall on the practice sequences (Desrochers et al., 2015, 2019; Schneider & Logan, 2006; Trach et al., 2021). Once this behavioral threshold was reached, participants were scanned while performing the task. The same equipment was used for training as for displaying the task and making responses during scanning. Stimuli were projected onto a 24” BOLDscreen 32 UHD and the task was run using Psychtoolbox on Matlab 2017b.

#### Data Acquisition

A Siemens 3T PRISMA MRI scanner with a 64-channel head coil was used for whole-brain imaging. Functional data for two of the 50 participants were acquired using an echo-planar imaging pulse sequence (repetition time, TR = 2.0 s; echo time, TE = 28 ms; flip angle 90°; 38 interleaved axial slices; 3.0 x 3.0 x 3.0 mm). Anatomical scans included a T1-MPRAGE (TR, 1900 ms; TE, 3.02 ms; flip angle, 9.0°; 160 sagittal slices; 1.0 x 1.0 x 1.0 mm) and a T1 in-plane scan (TR, 350 ms; TE 2.5 ms; flip angle, 70°; 38 transversal slices; 1.5 x 1.5 x 3.0 mm). The remaining 48 participants were scanned on an updated protocol designed to enhance signal to noise ratio of the data. Functional data for these participants were acquired using an echo-planar imaging pulse sequence (repetition time, TR = 1.53 s; echo time, TE = 33 ms; flip angle 62°; 60 interleaved axial slices; 2.4 x 2.4 x 2.4 mm). Anatomical scans included a T1-MPRAGE and a T1 in-plane scan with the same parameters as in the original protocol.

### Data Analysis

#### Behavior

All behavior analyses were conducted using custom scripts in Matlab 2023a. As in previous studies using the same or similar sequential tasks (Desrochers et al., 2015, 2019; Schneider & Logan, 2006; Trach et al., 2021), the following sets of trials were excluded from analyses. The first four trials (first sequence) in every block were removed across participants (approximately 1.6% of trials per participant) to prevent changes in reaction times (RTs) at block initiation from confounding with RT changes due to sequence initiation or task switching. Additionally, trials were excluded that had RTs < 100 ms (< 1 % of trials per participant) to prevent inclusion of trials in which categorization choices were guessed. Error rates (ERs) were calculated on the remaining trials. Periods of trials were also removed in which participants “lost track” of the sequence. These trials were defined as “lost” for 2 or more error trials up until the next 4 correct adjacent trials occurred (approximately 6.4% of trials per participant). “Lost” trials were excluded to ensure all analyzed trials were ones in which the participants were completing the task as instructed. Statistical analyses were conducted on RTs and ERs using RM-ANOVAs and t-tests.

Age was included as a covariate in ANOVAs that compared across groups to control for potential differences in age distribution across samples. Sequence initiation cost was calculated as the difference in position 1 and position 3 RTs across sequence types. This calculation averaged RTs across all trials for each participant by positions 1 and 3. These averaged RTs were subsequently subtracted, resulting in one initiation cost number per participant. Switch costs were defined as the RT and ER differences between switch and repeat trials. Switch costs were calculated by averaging RTs and ERs across all trials that are switches, which excluded position 1 and included positions 2 and 4 in complex sequences and position 3 in simple sequences and subtracting averaged RTs and ERs across all repeat trials, which excluded position 1 and included position 3 in complex and positions 2 and 4 in simple sequences. This calculation results in an average switch cost number per participant. ANOVAs testing behavior hypotheses examined elements of the costs (sequence positions 1 and 3 for initiation cost and switch and repeat trials for switch costs). We additionally report results of the subtraction that gives the behavior costs as well, and use these costs calculations for symptom severity correlations. Specifically, clinical symptom measures (Y-BOCS) were correlated (pairwise linear) with behavior costs and neural activity in OCD.

#### Preprocessing

All imaging data were preprocessed using Statistical Parametric Mapping (SPM12) in Matlab 2017b. Participants with motion exceeding one voxel (3.0 mm for the first two participants and 2.4 mm for the remaining 48 participants) were excluded from analysis. Images were then resampled to account for differences in acquisition timing and matched to the first slice. All images were then corrected for motion using B-spline interpolation and normalized to the Montreal Neurological Institute (MNI) stereotaxic template with affine regularization. Lastly, data were smoothed using an 8mm full-width at half-maximum Gaussian kernel and resampled using trilinear interpolation.

#### FMRI Models

All general linear models were constructed using SPM12 and custom scripts in Matlab 2023a. Onset and parametric regressors were convolved with the canonical hemodynamic response function (HRF). Additionally, onset regressors were convolved with the first time derivative of the HRF. Nuisance regressors were included to account for variance due to translational and rotational motion (x, y, z, roll, pitch, yaw) and for the first four trials (first sequence) of every block, time during instruction, and sequence position question trials.

Beta values related to regressors were estimated using a subject-specific fixed-effects model. Whole brain contrasts estimated subject-specific effects, and these estimates were entered into a second-level analysis with subject treated as a random effect. T-values resulting from these contrasts were used for analyses. Whole brain group voxel-wise effects were corrected for multiple comparisons using extent thresholds at the cluster level to yield family-wise error correction and were considered significant at P < 0.05. Group level contrasts were rendered on a 3D brain using Connectome Workbench (humanconnectome.org/software/connectome-workbench).

*<u>Onsets models:</u>* We constructed stimulus onset regressors to model univariate effects at each sequence position. These regressors were modeled as 0 second durations at the onset of each stimulus. Separate regressors were included for each position in the sequence (1-4) and each sequence type (complex and simple), for a total of eight regressors for the conditions of interest (**Figure 2A,C**).

**Figure 2.**
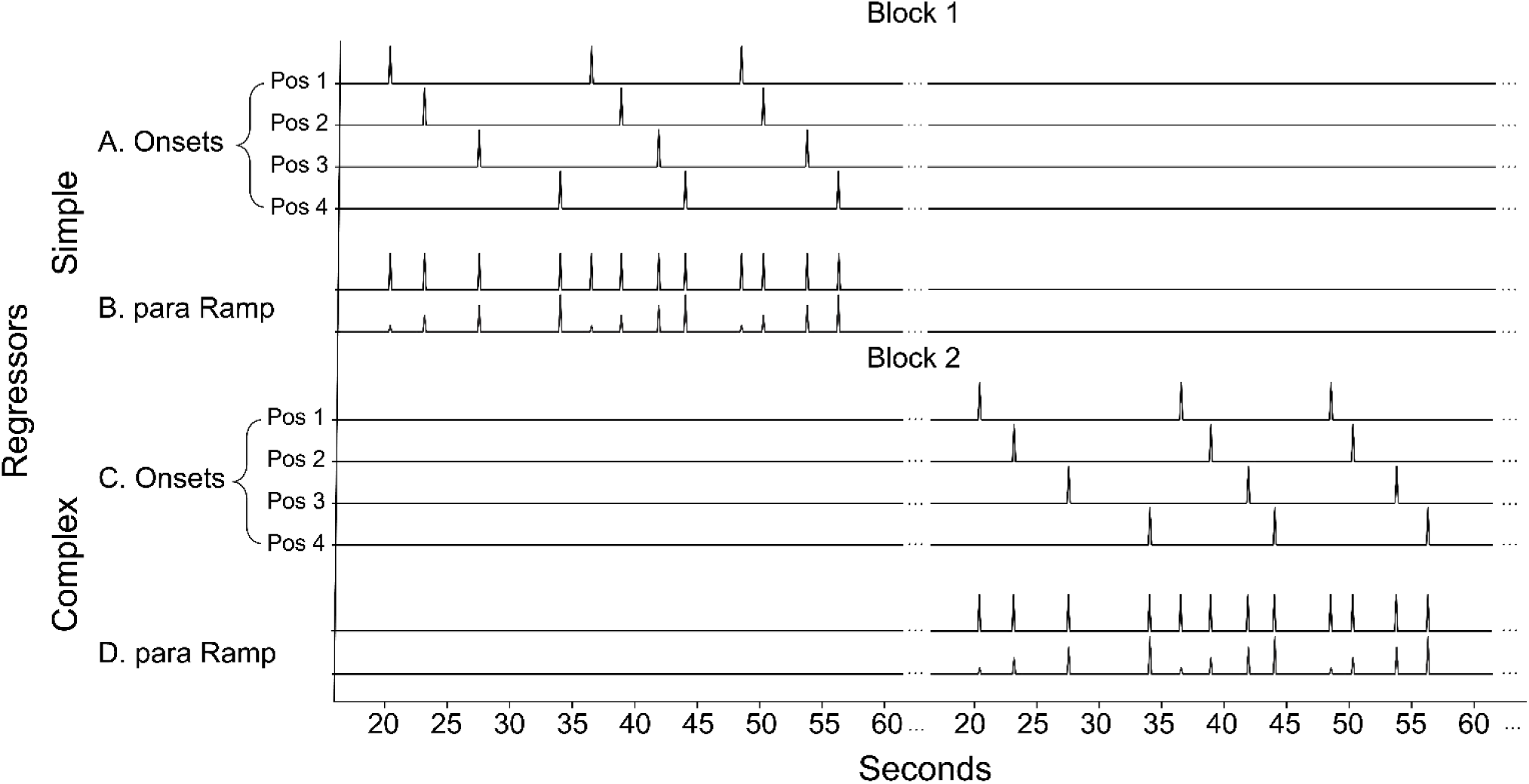
Illustration of unconvolved onsets and parametric regressors for simple and complex sequence blocks. **A.** Onset regressors are set for each sequence position (1-4), as a zero duration stick function. In this example, onset regressors are shown in a block that is composed only of simple sequences (pattern “AABB”), while no regressors are set for simple sequences in the second block, which is composed of only complex sequences (pattern “ABBA”). Each block only contains one sequence type (simple or complex) and the order of the blocks is counterbalanced. **B.** Parametric ramp regressors are composed as a monotonic increase 1-4 by sequence position, making a positive slope across positions. Onsets are modeled together with the parametric ramp regressor in the parametric model. In the parametric model, onsets are estimated first followed by the parametric ramp regressor, so that any variance assigned to the parametric ramp is above and beyond that attributed to the onset regressors. **C.** Onset regressors illustrated for the second block, composed of only complex sequences. **D.** Parametric ramp regressors illustrated for the second block, composed of only complex sequences.

*<u>Parametric ramp model</u>*: To test for ramping activity, we constructed a regressor for each sequence type (complex and simple) that included a zero-duration onset for each stimulus (**Figure 2A,C**) and a parametric (numbers 1-4) for a linear increase across the four positions in the sequence (**Figure 2B,D**). In the parametric model, onsets were modeled with the parametric ramp (**Figure 2B,D**) and estimated hierarchically. Onsets were estimated first, followed by the parametric ramp regressors, so that variance assigned to the parametric regressor was above and beyond what could be accounted for by the stimulus onset alone.

Each model was used to address different levels of control. Because, by definition, ramping spanned entire sequences, it was used to assess abstract sequence related activity throughout the task (All Ramp > Baseline) and by sequence type (Complex > Simple Ramp). It was not feasible to assess task switching using the ramp model, as task switches and repeats happen at individual trials within sequences. To directly examine neural activity related to the level of task sequential control, we examined onset activity for all switch trials compared to all repeat trials throughout the task, in the All Switch > Repeat and No Position 1 Switch > Repeat contrasts. The All Switch > Repeat contrast was used to assess onset activity related to all switches and repeats in the task across all conditions. The No Position 1 Switch > Repeat contrast excluded trials at position 1, to account for potential sequence initiation effects, thereby isolating within-sequence switches and repeats.

#### ROI Analysis

Region of interest (ROI) analyses complemented whole-brain analyses. ROIs for replication analyses were taken from a previous study (Desrochers et al., 2015). ROIs were defined from significant peaks of activation from the Onsets model voxelwise contrasts No Position 1 Switch > Repeat and Position 2,3 Switch > Position 2,3 Repeat. These contrasts omitted the first sequence position to ensure only trial-based switching and repeating were being captured. ROIs were also defined from significant activation peaks from the Parametric model contrast Parametric Ramp > Baseline, as constructed in Desrochers et al., (2015). We extracted T values from these ROIs using these contrasts. Repeated measures analysis of variance (RM-ANOVAs) or t-tests were subsequently performed on these values.

## Results

### OCD participants exhibit differences in task level behavior

To address questions of potential behavioral and neural deficits in abstract sequential control and task switching in OCD, two groups of participants (OCD and healthy control, HC) completed abstract cognitive task sequences (**Figure 1**) while undergoing fMRI scanning. Briefly, participants were presented at each block start with four-item sequences of simple categorization decisions, either simple (containing one task switch, e.g. shape, shape, color, color) or complex (containing two task switches, e.g., shape, color, color, shape). On each trial, participants used information about sequence position to correctly categorize the color or shape of the image. Participants repeated sequences until the end of each block. To probe neural mechanisms underlying sequential behavior, participants completed five runs, each containing four blocks of this task while undergoing fMRI scanning. Two features of this task are relevant to assessing performance: a feature at the level of abstract sequential control (initiation cost) and a more general cognitive control feature at the task level of sequential control (switch cost) (Desrochers et al., 2015; Schneider & Logan, 2006). Initiation cost is the difference in reaction times (RTs) between sequence positions 1 and 3 (both positions are repeats or switches, to account for trial type effects), while switch cost (Monsell, 2003) is the RT or ER difference between switch and repeat trials, excluding the first position.

We first examined behavioral markers of sequential and task level control separately in each group to replicate previous results and show that participants were performing the task as instructed. Participants in both HC and OCD groups replicated sequential and cognitive control effects observed previously. Overall, participants in both groups performed well (RT means +/- 1 SD: HC, 1.23 +/- 0.29 s; OCD: 1.32 +/- 0.29 s; ER means +/- 1 SD: HC: 7.78 +/- 7.33; OCD: 8.52 +/- 7.04). To assess sequence level control, we examined initiation costs by comparing RTs at the first and third positions of the sequence, which compares the start and subsequent position in a sequence, holding the trial type constant (e.g., in complex sequences, position 1 and 3 are both task repeats). Initiation costs were examined specifically in RTs because this effect has been previously observed primarily in RT in sequential tasks (Desrochers et al., 2015, 2019; Schneider & Logan, 2006; Trach et al., 2021). We found a significant difference between position 1 and 3 in both groups (one-way RMANOVAs: HC: F(1,24) = 58.50, p < 0.001, ηp^2^ = 0.71; OCD: F(1,24) = 44.90, p < 0.001, ηp^2^ = 0.65; initiation cost means +/- 1 SD: HC, 0.15 +/- 0.09 s; OCD, 0.18 +/- 0.13 s). Further, we found that initiation costs were significantly greater in complex compared to simple sequences in both groups (initiation cost means +/- 1 SD: HC complex: 0.23 +/- 0.15 s; simple: 0.08 +/- 0.12 s; t(48) = 3.89, p < 0.001; OCD complex: 0.24 +/- 0.18 s; simple: 0.12 +/- 0.13 s; t(48) = 2.79, p = 0.01). These results replicate previous studies in healthy participants (Desrochers et al., 2015; Schneider & Logan, 2006), indicating that participants were adhering to instructed sequence boundaries, i.e. not simply alternating between two of one task and two of the other. At the task control level, we compared task switches to task repeats in RT and ER. We found significant differences between switches and repeats for both groups in RTs (one-way RMANOVAs: HC: F(1,24) = 17.60, p < 0.001, ηp^2^ = 0.42; OCD: F(1,24) = 27.50, p < 0.001, ηp^2^ = 0.53; switch cost means +/- 1 SD: HC, 0.13 +/- 12 s; OCD, 0.11 +/- 0.08 s) but in ERs observed significant switch costs only in the HC group (HC: F(1,24) = 26.73, p < 0.001, ηp^2^ = 0.53; OCD: F(1,24) = 0.85, p = 0.36, ηp^2^ = 0.03; switch cost means +/- 1 SD: HC, 1.29 +/- 1.25; OCD: 0.35 +/- 1.89). These results replicate task level control behavioral markers for switch costs in RTs (Desrochers et al., 2015; Rogers & Monsell, 1995; Schneider & Logan, 2006). Taken together, these results show that both HC and OCD participants performed the task as instructed and exhibited behavioral markers of sequence and task level control.

To test for behavioral deficits in OCD at the sequence and task levels of control, we used the same measures to compare RTs and ERs to HCs. For sequence level control, we found that sequence initiation (position 1 vs. 3) RTs were not different across the groups (**Table 2**, **Figure 3A**). At the task level of control, task switching RTs were not different between the groups (**Table 3**, **Figure 3A**). However, ERs were greater for repeat trials in participants with OCD resulting in smaller switch costs (group x trial type **Table 3**, **Figure 3B,C**). These behavioral results suggest that abstract sequential control was preserved while task level control was different and possibly impaired during an abstract sequential paradigm in OCD.

**Figure 3.**
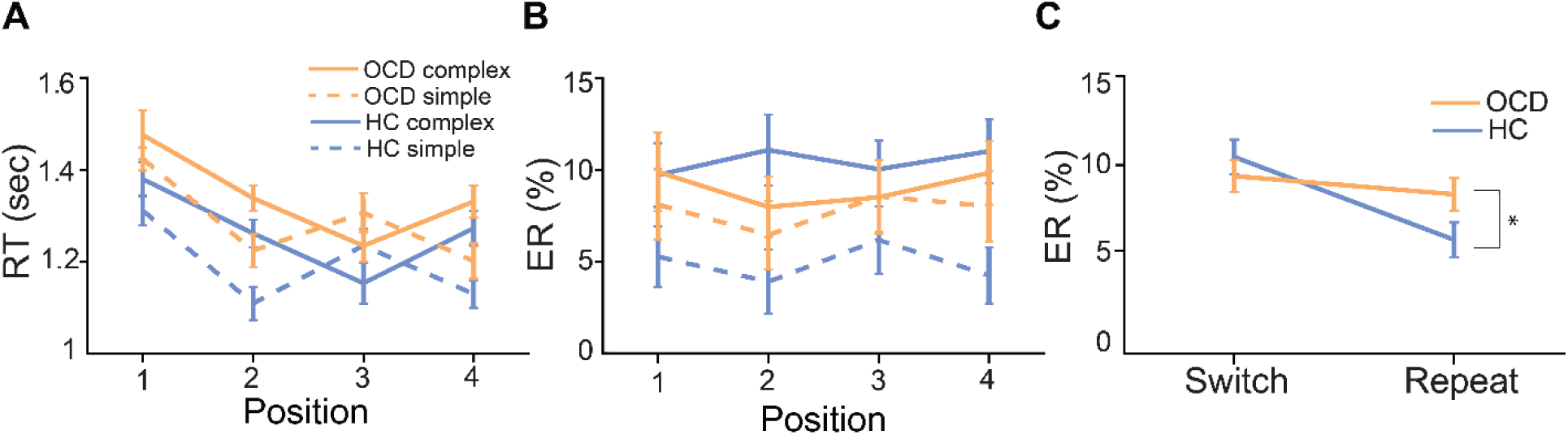
ER task switching deficits occur in OCD compared to HCs. **A**. RTs between HCs and OCD do not significantly differ across simple and complex sequences. **B**. ERs significantly differ between HCs and OCD across sequence positions. **C**. There is a significant interaction in ERs by trial type (switch vs. repeat trials) between groups (indicated by “*”).

**Table 2.** RM ANOVAs of RTs (s) to assess initiation costs between groups. Dfs, F statistics, P values, and effect sizes (ηp2) are reported in each column. To probe initiation cost differences between groups we conducted an ANOVA with group, age, and position (1 and 3) as factors. We did not observe a significant interaction between group and position, indicating initiation costs were not significantly different between groups.

|  |  | Reaction Times (RTs) |  |  |
| --- | --- | --- | --- | --- |
| *Factor* | *dfs* | *F* | *p* | *eta2p* |
| Group | 1,47 | 0.70 | 0.40 | 0.02 |
| Age | 1,47 | 1.20 | 0.27 | 0.03 |
| Position (1 vs. 3) | 1,47 | 8.20 | 0.01 | 0.15 |
| Age x Position | 1,47 | 0.10 | 0.71 | 0.00 |
| Group x Position | 1,47 | 1.00 | 0.32 | 0.02 |

**Table 3.** RM ANOVAs of RTs (s) and ERs (%) to assess switch costs between groups. Dfs, F statistics, P values, and effect sizes (ηp2) are reported in each column. To probe switch cost differences between groups we conducted an ANOVA with group, age, and trial type (switches and repeat trials) as factors. We did not observe a significant group by trial type interaction in RTs, but we did observe a significant interaction in ERs.

|  |  | Reaction Times (RTs) |  |  | Error Rates (ERs) |  |  |
| --- | --- | --- | --- | --- | --- | --- | --- |
| *Factor* | *dfs* | *F* | *p* | *eta2p* | *F* | *p* | *eta2p* |
| Group | 1,47 | 0.7 | 0.41 | 0.01 | 0.1 | 0.8 | 0 |
| Age | 1,47 | 1.40 | 0.24 | 0.03 | 0.4 | 0.55 | 0.01 |
| Trial type (switch vs. repeat) | 1,47 | 11.6 | 0.00 | 0.20 | 2.4 | 0.13 | 0.05 |
| Age x Trial type | 1,47 | 1.9 | 0.18 | 0.04 | 0.2 | 0.7 | 0 |
| Group x Trial type | 1,47 | 0.6 | 0.44 | 0.01 | 4.4 | 0.04 | 0.09 |

Because we observed behavior deficits in ERs in the current study, we next tested if the behavior differences in OCD correlated with symptom severity. Specifically, we assessed if and how ER switch costs correlated with OCD symptom severity (total Y-BOCS scores). This hypothesis was motivated by a previous study that reported a positive correlation between OCD symptom severity and deficits in cognitive control (Remijnse et al., 2013). We observed a marginally positive correlation between ER switch cost and symptom severity scores (Y-BOCS p = 0.06, r = 0.38), such that OCD participants with higher symptom severity scores exhibited marginally greater switch costs.

### Neural indicators of sequence and task levels of control replicate previous studies

The present behavioral task was used in a previous neuroimaging study in HCs (Desrochers et al., 2015); therefore, we first replicated neural effects related to sequence and task level control. We examined a specific neural dynamic, increasing activation across items in each sequence (“ramping”), as an indicator of sequential control. To initially examine ramping dynamics in this population of participants, we first aimed to replicate the existence of a distribution of brain areas that show this dynamic during the task. Ramping dynamics have been robustly associated with a variety of sequential tasks (Desrochers et al., 2015, 2019; McKim & Desrochers, 2022). Ramping was modeled as a parametric increase in BOLD activation across the four positions of each sequence (i.e., resetting at position 1) that explained variance above and beyond stimulus onsets (**Figure 2**). Though we had hypotheses about the involvement of specific regions in this task (i.e., the RLPFC), we first wanted to establish the general presence of ramping activation. To test for this activity, we created a single large ROI that contained all the significant ramping clusters from the All Parametric > Baseline contrast in (Desrochers et al., 2015). We found significant ramping activity in this combined ROI in each group separately (OCD: t(24) = 2.98, p = 0.001, HC: t(24) = 3.39, p = 0.002), and no difference in ramping between groups (t(48) = −0.14, p = 0.88), replicating neural effects of sequential control in OCD and HC groups in this study.

We next measured neural responses to task switching as an indicator of task level control (Monsell, 2003). To test for neural activity related to task switching, we created ROIs from regions previously observed to have significant Switch > Repeat neural activity (Desrochers et al., 2015) (see Methods). We found significant or marginal activity across all participants in the majority of ROIs in these conditions (L occipital: t(49) = 3.16, p < 0.001, R IFG [No Position 1 Switch > Repeat]: t(49) = 1.83, p = 0.05, R SMA/cingulate: t(49) = 1.71, p = 0.06, R IFG [Position 23 Switch > Position 23 Repeat]: t(49) = 2.03, p = 0.04). Activity related to task switching was not significantly different between OCD and HCs in any of the ROIs (t(48) = −1.02, p = 0.5, all ROIs combined), replicating neural responses to task switching.

### Abstract sequential control related ramping differs in OCD and HC

To investigate if participants with OCD showed neural activity differences related to abstract sequential control, we focused on three sets of analyses. First, we compared activity in the RLPFC between groups due to this region’s necessity for abstract sequence performance (Desrochers et al., 2015). Second, we examined ramping dynamics across sequences in the whole brain to determine differences in the network of areas engaged in sequence-related dynamics. Third, to determine if there were differences based on the demands of each sequence type, we compared ramping activity between complex and simple sequences.

We first tested the hypothesis that there is decreased activity in RLPFC in participants with OCD compared to HC. Although we observed no sequential behavior deficits in OCD (**Figure 3**), it is possible differential RLPFC activity occurs in OCD participants compared to HCs. To test this hypothesis, we used an ROI defined by the RLPFC cluster of ramping activation in a previous study (Desrochers et al., 2015), hereafter referred to as the D15 ROI. D15 is causally involved in performing abstract sequences, as transcranial magnetic stimulation (TMS) to this region selectively produced task deficits in HCs in the same task as used in the present study (Desrochers et al., 2015). In the RLPFC D15 ROI, there was no significant ramping difference between groups (**Figure 4A**; All Parametric Ramp > Baseline contrast; t(48) = −0.36, p = 0.72). Acknowledging that group differences in activity in the RLPFC may not be limited to ramping dynamics, and because hypoactivation has been observed in cognitive control related areas of the lateral frontal cortex in OCD (Gu et al., 2008; Remijnse et al., 2013), we also examined onset activity in the D15 ROI. We found no significant difference in overall (onset) activity between the groups (**Figure 4B**; All > Baseline contrast; t(48) = −1.12, p = 0.27). Therefore, we did not observe RLPFC differences between the groups, possibly supporting behavioral findings that there were no abstract sequential control differences between OCD and HCs.

**Figure 4.**
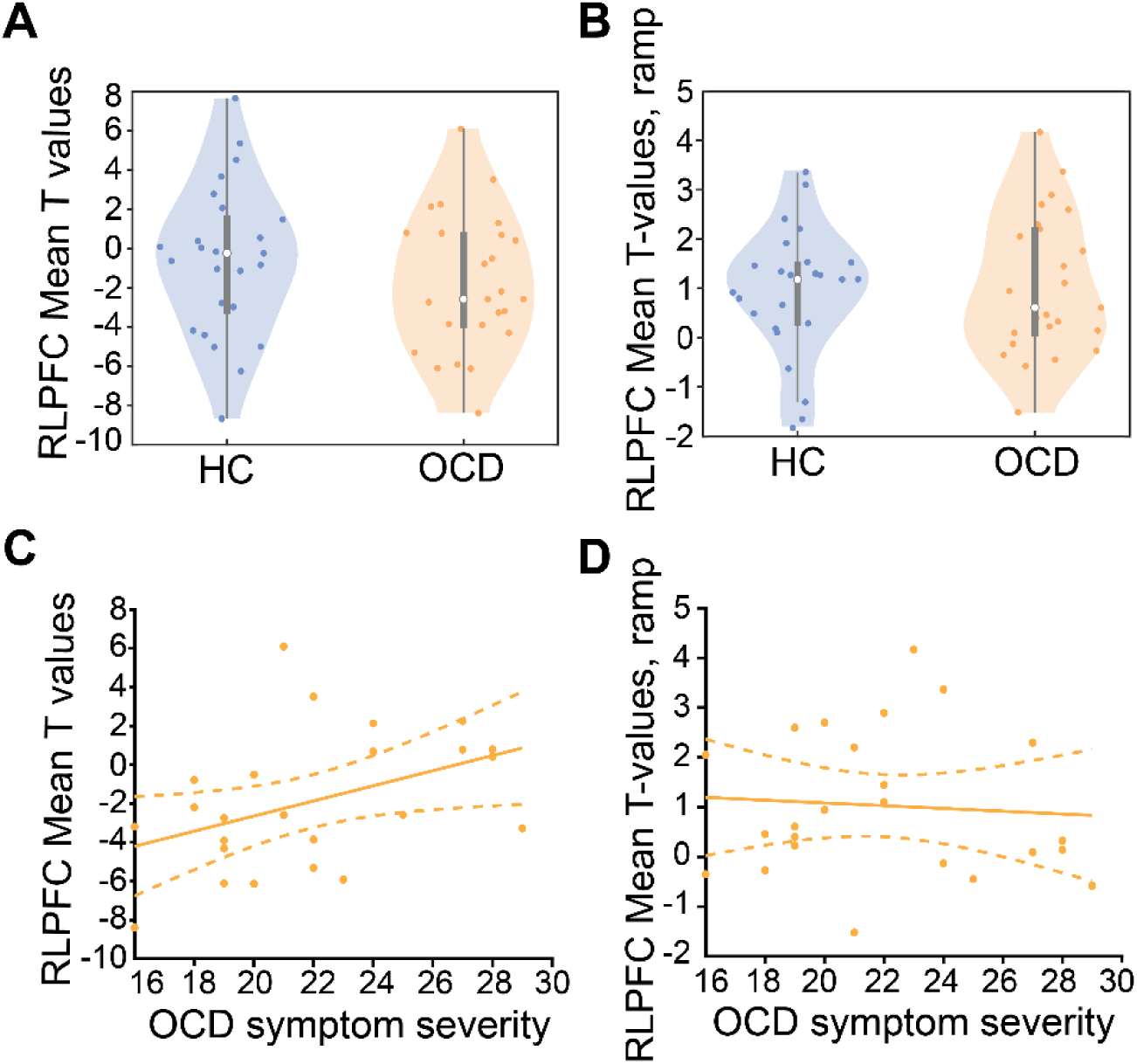
RLPFC ramping and onset dynamics do not differ between groups but activity correlates with OCD symptom severity. **A**. Ramping activity in the D15 ROI does not differ between HCs and OCD **B**. All > Baseline activity in the RLPFC ROI does not differ between HCs and OCD. **C.** OCD symptom severity (total Y-BOCS scores) does not significantly correlate with D15 ROI ramping in OCD. **D**. OCD symptom severity (total Y-BOCS scores) positively correlated with All > Baseline activity in the RLPFC.

Although there were no differences in ramping or overall activity in the RLFPC between groups, to gain a more complete picture of any potential interaction between neural activity and OCD we correlated RLPFC activity with OCD symptom severity.

There was no correlation between RLPFC ramping and symptom severity (**Figure 4C**; r = −0.03, p = 0.90), providing further support that RLPFC function and abstract sequential control as measured by ramping activation is not impaired in OCD. However, we did find that OCD participants with greater symptom severity (total Y-BOCS score) showed greater overall (onset) task activity in the RLPFC resulting in a significant positive correlation (**Figure 4D**; p = 0.037, r = 0.43). These results suggest that although onset activity does not significantly differ between groups, OCD symptom severity may still influence activity levels in the RLPFC, a region crucial for abstract sequence execution (see Discussion).

To follow up on ROI analyses, we next tested whole brain activity related to abstract sequential control. We examined ramping dynamics across and between sequence types (complex and simple) to determine if there were differences in the network of areas that support abstract sequence performance between OCD and HC participants. Motivated by previous studies showing differences between OCD and HC in cognitive control areas, we hypothesized that areas outside the RLPFC may be differentially involved in abstract sequence control in OCD to produce behavior that looks similar to HCs.

Whole brain ramping activity supported ROI results and revealed brain areas that were significantly different between OCD and HC. First, significant RLPFC ramping (All Parametric Ramp > Baseline) was observed in HC (**Figure 5A**) and OCD (**Figure 5B**) separately, supporting the D15 ROI results. Further, significant ramping activation was observed across a common group of frontal, parietal, and dorsal medial cortical areas (**Table 5**). However, when directly contrasting ramping activity between OCD and HC, significantly greater ramping in OCD was observed in the pregenual anterior cingulate cortex (rACC) and posterior superior frontal sulcus (SFS), a region near the posterior DLPFC, and posterior supplementary motor area (pSMA/SMA) (**Figure 5C**; **Table 5**).

**Figure 5.**
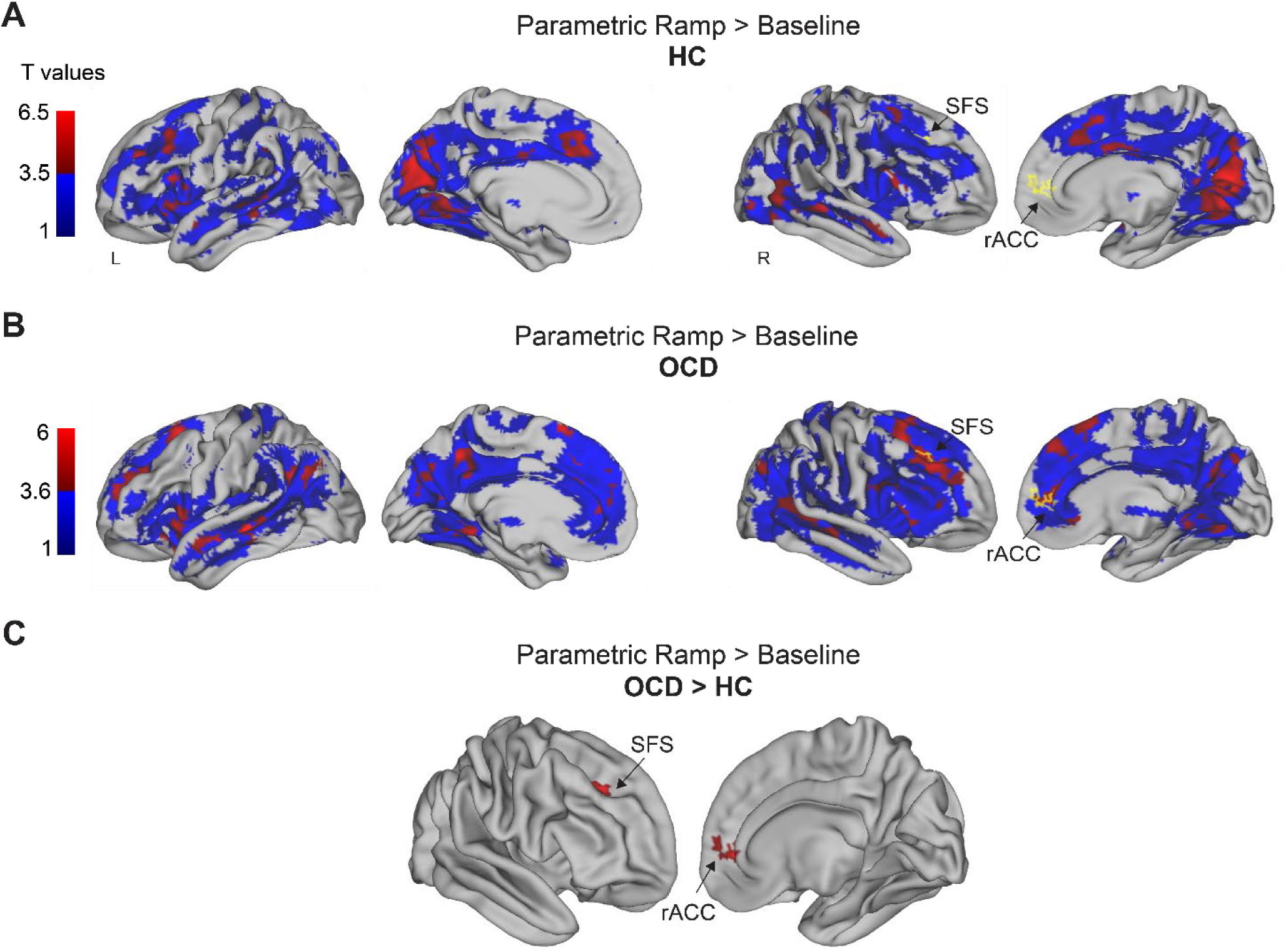
Ramping activity in the SFS and rACC dissociates OCD from HCs during abstract sequential control. **A**. Whole brain contrast Parametric Ramp > Baseline in HCs. Sub threshold activity (p < 0.05 uncorrected) is in blue and activity FWE cluster corrected at p < 0.05, height p < 0.005, extent 167 voxels is shown in red. Color bar denotes increasing T values, with dark to light blue reflecting increasing activity below threshold, and dark red to light red reflecting increasing activity that is above threshold. Activity that significantly dissociates OCD from HC (shown in [C]) is outlined in yellow for reference. **B**. Whole brain contrast Parametric Ramp > Baseline in OCD. Sub threshold activity (p < 0.05 uncorrected) is in blue and activity shown in red is FWE cluster corrected at p < 0.05, height p < 0.005, extent 167 voxels. Color bar reflects increasing T values in the same manner as in (A). Activity that significantly dissociates OCD from HC (shown in [C]) is outlined in yellow for reference. **C.** Parametric Ramp > Baseline, FWE cluster corrected at p < 0.05, height p < 0.005, extent 822 voxels, OCD > HC, ramping activity that is present in OCD but not in HCs. SFS: superior frontal sulcus, rACC: rostral anterior cingulate cortex.

**Table 5.**
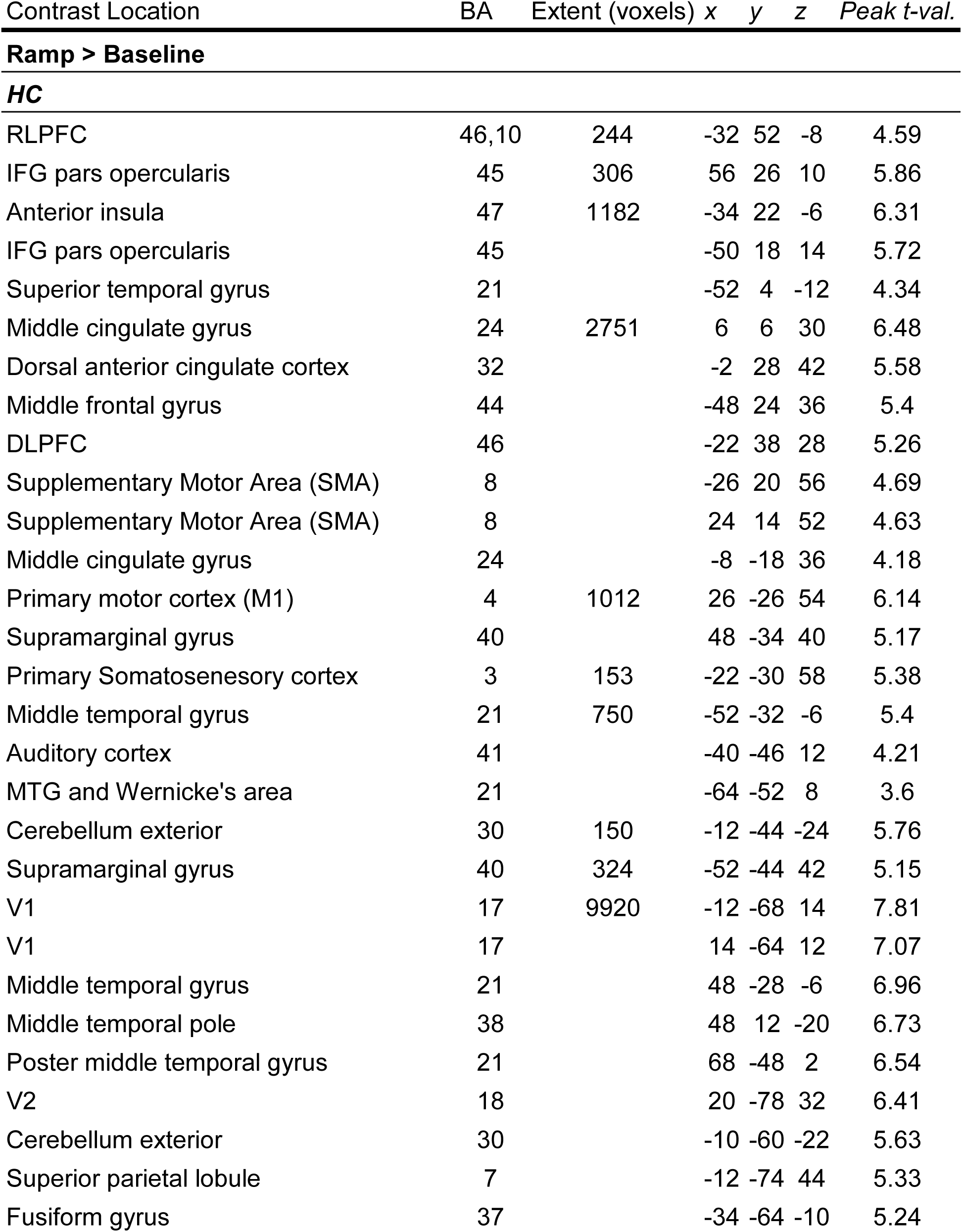
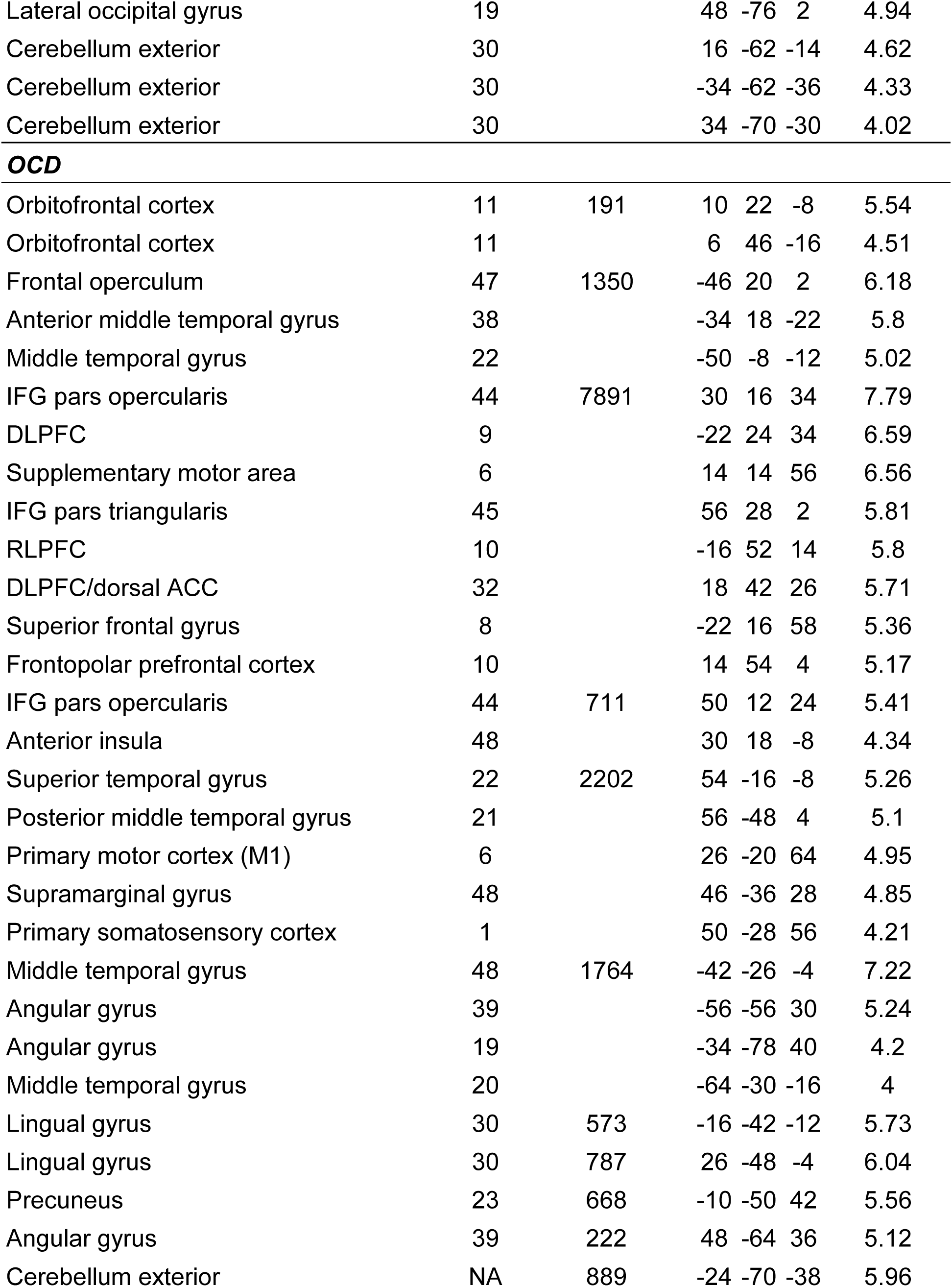
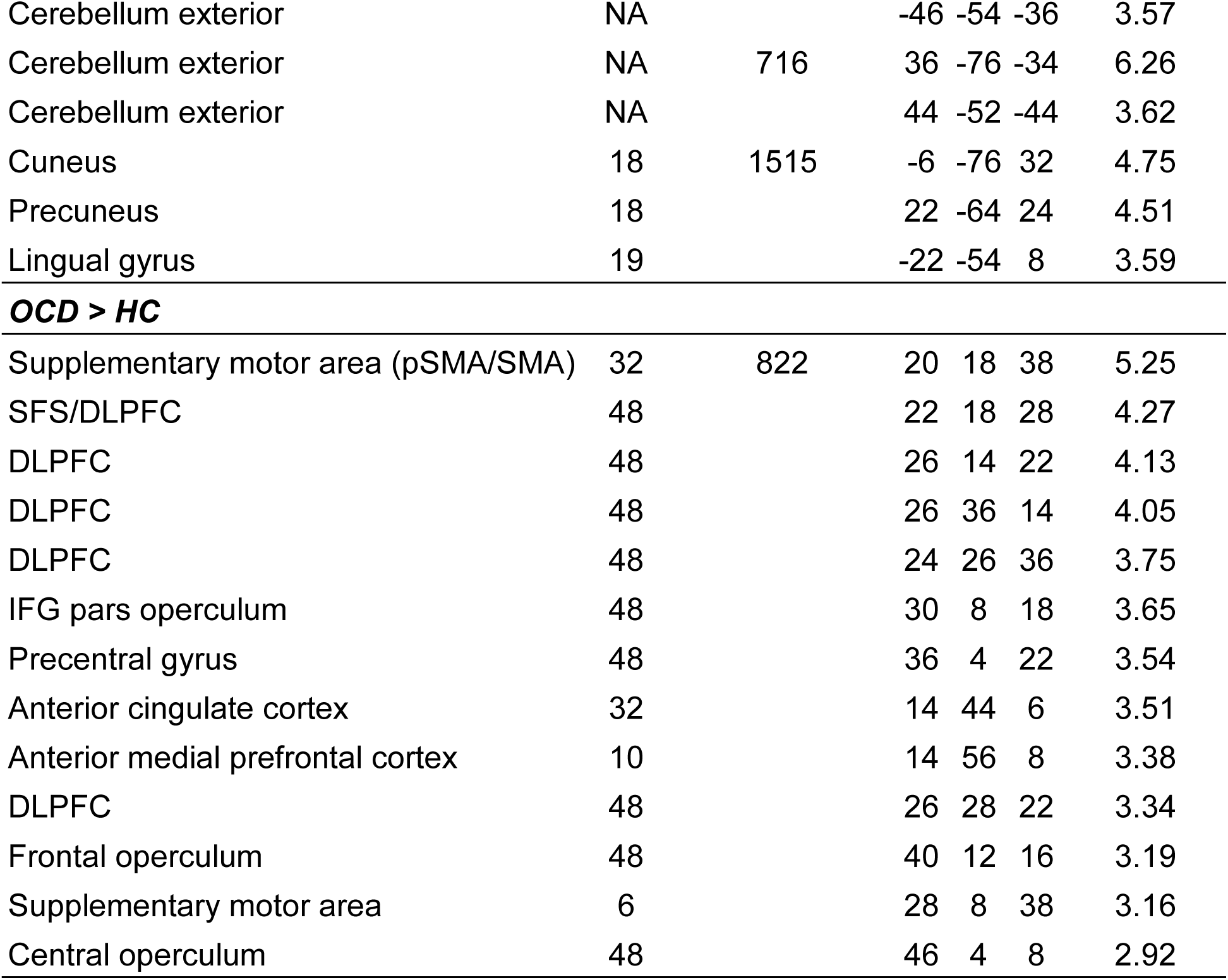
Activation coordinates, significant ramping activity in the Ramp > Baseline contrast in HC, OCD, and OCD > HC. Clusters reliable at p < 0.05 corrected. No clusters in the HC > OCD Ramp > Baseline contrast survived correction. Extent p < 0.001 for the OCD and HC contrasts, and p < 0.005 for the OCD > HC contrast. Distance between significant clusters was set to 25 mm for the HC and OCD contrasts. Distance between significant clusters was set to 12 mm for the OCD > HC contrast. Coordinates are the center of mass in MNI.

We note that subthreshold activation is included solely for illustrative purposes in the individual groups (**Figure 5 A,B**) to aid in interpreting the limited number of areas that are statistically different in these direct comparisons. No clusters of activation survived correction in the reverse, HC > OCD, ramping contrast. Because these greater activity in the rACC and SFS are defined by a dynamic that spans entire sequences (ramping) they are sequence related. Therefore, these results provide partial evidence for the hypothesis that abstract sequential control differs in OCD and HCs. We will return to the potential role of these areas in abstract sequences in the Discussion.

Next, we tested ramping dynamics in different sequence types related to the level of abstract sequential control. First, this analysis serves as a control, as we would not expect ramping that differs by sequence type to occur in the RLPFC. Previous studies established that this dynamic generalizes across different sequence variations (Desrochers et al., 2015, 2019). Second, we do expect ramping to occur differentially across sequence types in other cortical regions outside of the RLPFC (Desrochers et al., 2015). Because, by definition, these dynamics extend through entire sequences, such ramping differences may be related to increasing cognitive control resources needed for complex versus simple sequences and to the abstract sequence control level. Based on literature showing dysfunctional cognitive control in OCD (Gu et al., 2008; Remijnse et al., 2013), ramping in this contrast may differ in our OCD participants, in that there may be decreased ramping in complex versus simple sequences in these participants compared to HCs.

To test the hypothesis that ramping may occur differentially both by sequence type and by participant group, we tested group differences throughout the whole brain by examining activity in the Complex > Simple Parametric Ramp contrast. In this contrast we did not observe ramping differences in the RLPFC, but we did observe significantly increased ramping in HCs compared to OCD in a region of the medial temporal cortex (MTG), the supplementary motor area (SMA), and the temporal occipital junction (TOJ) (**Table 6**). There were no clusters that reached statistical significance in the reverse OCD > HC contrast. We observed the same clusters of increased ramping in OCD > HC in the Simple > Complex Ramp contrast (**Figure 6A**; **Table 6**). The activity in these contrasts between groups was therefore an interaction of ramping by sequence type between groups in all three clusters (FWE corrected at p < 0.05, extent p < 0.005). Such an interaction could be produced by many different relationships among complex sequences, simple sequences, OCD, and HC. Therefore, to visualize this interaction we examined the T-values in each of the three clusters. All three areas showed a similar pattern of activity, with HC showing greater ramping in complex than simple sequences and OCD showing nearly the reverse pattern with greater ramping in simple than complex sequences (**Figure 6B,C,D**). We emphasize that these clusters were defined by the contrasts (and therefore are biased) and are solely for visualization purposes (confirmatory sequence type x group for clusters; SMA: F(1,48) = 8.58, p = 0.01; MTG: F(1,48) = 19.48, p < 0.001; TOJ: F(1,48) = 10.72, p = 0.002). These ramping results overall show that participants with OCD recruit additional cortical regions to support sequential behavior compared to HCs.

**Figure 6.**
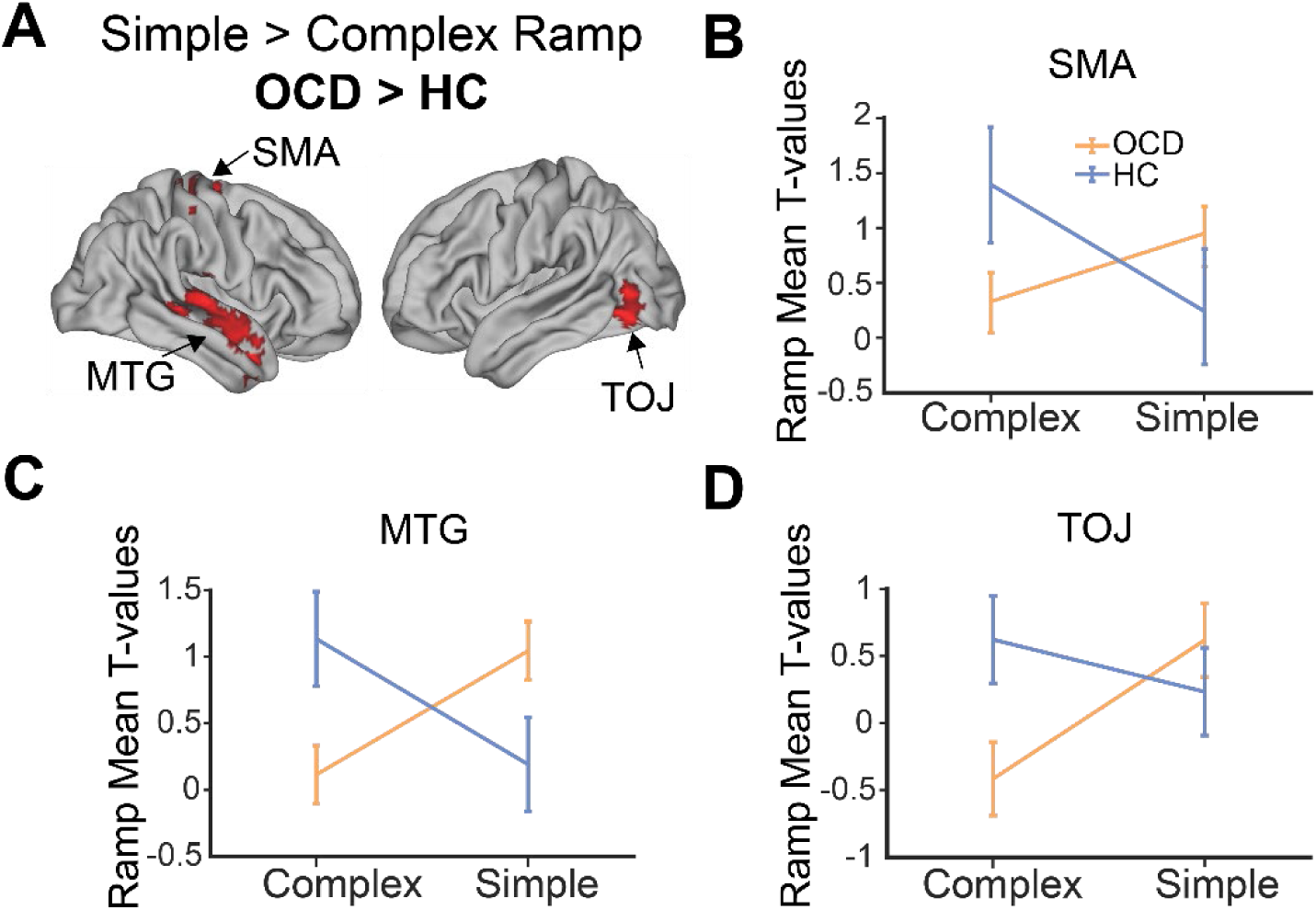
Ramping activity in novel cortical regions dissociates OCD from HCs. **A**. Whole brain contrast Simple > Complex Ramp, OCD > HC (FWE cluster corrected at p < 0.05, height p < 0.005, extent 330 voxels). **B.** Ramping in OCD vs. HCs produces significant interactions in the SMA, such that there is significantly more ramping in this region in Complex > Simple sequences in HC > OCD and significantly more ramping in Simple > Complex sequences in OCD > HC. **C.** Same as (**B**) but in the MTG. **D.** Same as (**B**) and (**C**) but in the TOJ. Biased ROIs are used for analyses in **B-D** purely for illustrative purposes. SMA: supplementary motor area, MTG: medial temporal gyrus, TOJ: temporal occipital junction.

**Table 6.**
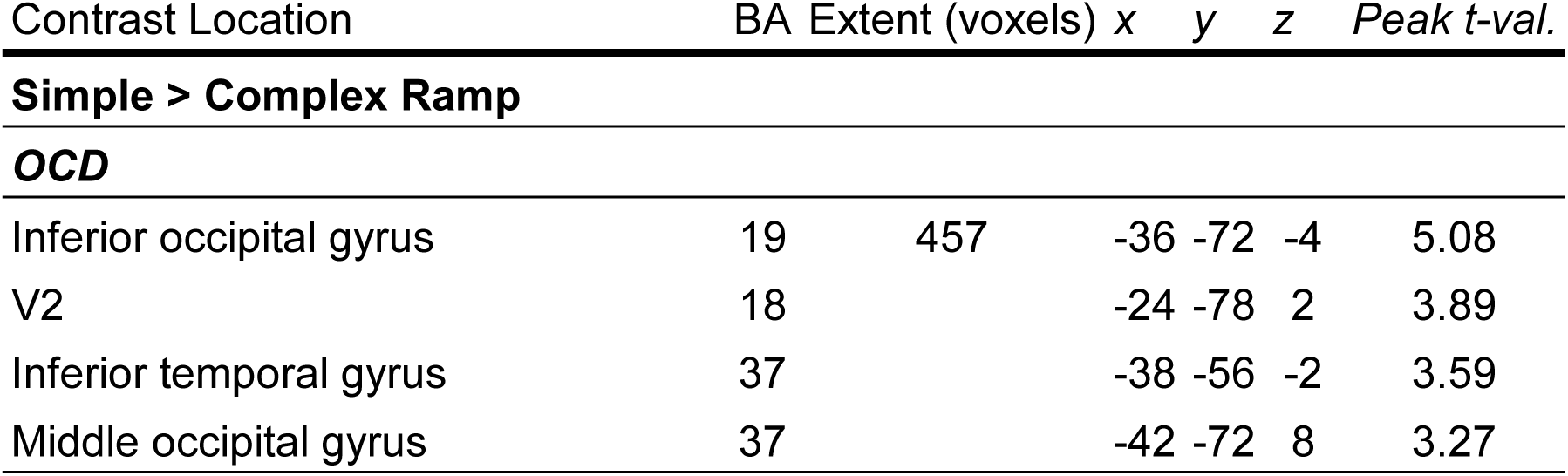
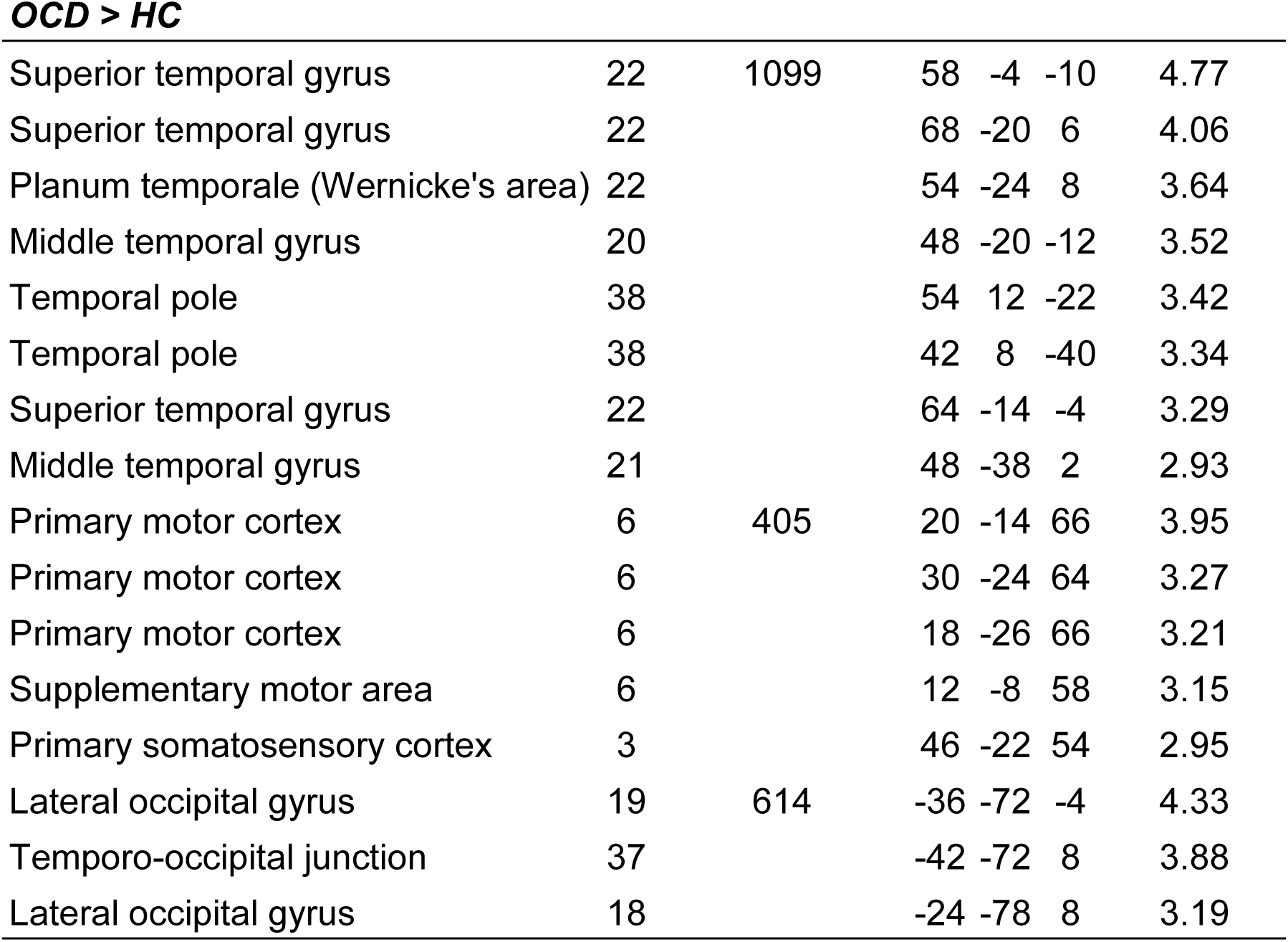
Activation coordinates, significant ramping activity in the Simple > Complex, Ramp contrast in OCD and OCD > HC. Clusters reliable at p < 0.05 corrected. No clusters in the HC and HC > OCD Simple > Complex Ramp contrast survived correction. Clusters in the HC > OCD Complex > Simple Ramp were the exact same as those in the OCD > HC Simple > Complex Ramp and were not reported for simplicity. Extent p < 0.005 was used for both the OCD and the OCD > HC contrasts. Distance between significant clusters was set to 12 mm. Coordinates are the center of mass in MNI. Extent p < 0.001 for the OCD and HC contrasts.

As an exploratory analysis, we correlated symptom severity with ramping in each of these clusters to complete hypothesis testing. We did not observe significant correlations between ramping activity in any of the observed regions and OCD symptom severity (rACC/SFS: r = 0.12, p = 0.61; SMA: r = 0.03, p = 0.87; MTG: r = 0.06, p = 0.76; TOJ: r = 0.19, p = 0.34). Therefore, ramping in these regions does not directly relate to symptom severity in the OCD participants.

### Task control level related activity does not dissociate OCD from HC

After testing hypotheses related to the level of abstract sequential control, we next tested the task level control. Specifically, we tested the hypothesis that compared to HCs, OCD participants would exhibit decreased activity in cortical regions associated with task switching. To test this hypothesis, we examined activity in the whole brain in the All Switch > Repeat and No Number 1 Switch > Repeat contrasts in the onsets model. This second contrast was included to control for any potential differences at position 1 of the sequence, given that RTs at this position are not reflective of task switching or repeating, but of sequence initiation. In both contrasts, no clusters of activation survived statistical correction in OCD > HCs or in HCs > OCD. Overall, these results show that neural activity related to task switching is not significantly different between groups.

## Discussion

We investigated the neural correlates of abstract sequence and task level control in OCD using fMRI. We hypothesized that in OCD abstract sequential control deficits would be associated with ramping and activity differences in the RLPFC along with differences in the broader network of areas that display ramping, and task switching deficits would be associated with decreased activity across cortical regions associated with this process. We found partial support for our first hypothesis such that abstract sequence control behavior and RLPFC activity (ramping and onsets) were not significantly different between the groups, but OCD participants showed significant differences in ramping in the rACC and SFS across sequence types and in the SMA, TOJ and MTG between sequence types (complex vs. simple). We also found partial support for our second hypothesis, in that behavioral switch costs were different in OCD, but without accompanying neural differences between the groups. In summary, these results suggest a group of brain areas, including some not previously identified as being associated with OCD such as the TOJ and MTG, differentially support abstract sequential control and related processes in OCD that cannot be explained by differences in task level control.

Ramping dynamics observed in the SFS and rACC suggest these regions are recruited more by OCD participants than by HCs to support cognitive control during abstract sequence completion. Previous work in HCs highlights ramping dynamics in both these regions related to cognitive control processes. For example, low frequency oscillations in the DLPFC increase across time when delaying responses during response inhibition (Khan et al., 2024), a control process utilized during task switching. Further, ramping in the ACC has been observed during related control processes that are invoked during sequencing, such as committed decision-making (Blanchard et al., 2015) and in preparation of switching task sets (Hyafil et al., 2009). Therefore, increased SFS/rACC ramping may signal the need in OCD to recruit additional control processes to complete the task. Additionally, ACC ramping occurs in anticipation of errors in anxiety-inducing conditions (Etkin et al., 2011), so increased rACC ramping may indicate increased control in response to anxiety provocation in OCD. Although both regions have been previously implicated as part of dysfunctional cortical circuits (McGovern & Sheth, 2017; Shephard et al., 2021), our work further highlights the importance of these regions in OCD, showing that they are recruited more heavily in these participants to complete sequences that require several types of cognitive control.

Increased ramping in other cortical regions (i.e., outside the RLPFC) suggests novel areas are recruited more in OCD than in HCs during the execution of simpler tasks, potentially engaging control processes invoked during abstract sequencing. Specifically, we showed OCD participants recruit cortical regions (MTG, TOJ, and the SMA) more than HCs during simple compared to complex sequences. Ramping in these cortical regions could support control processes invoked during abstract sequencing, as the SMA is recruited during motor sequencing (Tanji, 2001) and action monitoring (Bonini et al., 2014), the posterior MTG is activated during working memory, relevant to maintaining sequential information across time (Davey et al., 2016), and the TOJ has been shown to process the spatial frequency of visual stimuli (Kauffmann et al., 2014). Increased ramping in these regions may signal higher recruitment of sequential control-related processes is needed in OCD participants to execute simple sequences, whereas this additional recruitment in needed more in HCs during complex sequential control. This finding additionally may suggest an interaction between sequence and task level control, as all three of these areas support task switching in HCs (Chen et al., 2010; Timofeeva et al., 2024; Tsumura et al., 2021). While interactions between sequence and task level behavior have been shown in HCs (Trach et al., 2021), further studies would be necessary to evaluate this possibility in OCD. Overall, these findings suggest new cortical regions recruited more significantly in OCD participants to support abstract sequential task behavior.

Although we did not observe group differences in RLPFC dynamics or activity, the RLPFC may still be implicated in dysfunctional circuitry during task switching and sequential tracking in OCD during more anxiety-provoking task conditions. Although we did not directly investigate anxiety in the present study, a previous behavior-only study using the same abstract sequential control task reported sequence initiation deficits in participants with anxiety disorders but not in those with OCD, compared to HCs (Doyle et al., 2024), suggesting abstract sequential performance is weakened in anxiety disorders. In the present study, OCD symptom severity, which often co-occurs with anxiety co-morbidities, correlated with RLPFC activity during the task (**Figure 6C**).

Other work similarly shows anxiety correlates with activity in regions of the cognitive control network (Zhao et al., 2024) and specifically the RLPFC (Kaldewaij et al., 2021). Further, the lateral frontal pole encodes action-goal representations that are modulated by emotional valence in a task (Lapate et al., 2022). One compelling study showed that highly anxious individuals shift from utilizing the lateral frontal pole (including the RLPFC) to using the DLPFC and ACC to complete an emotionally valent action control task (Bramson et al., 2023). Taken together, the correlation between anxiety and RLPFC activity in the present study and these studies suggest that in OCD, the RLPFC may be underutilized in the sequential task specifically during conditions that provoke anxiety or are emotionally valent. Such conditions could provoke increased anxiety in this participant group, resulting in decreased activity or ramping in the RLPFC in conjunction with the observed increased ramping in the DLPFC and ACC. Such a scenario could also more closely imitate real-life OCD symptoms that produce anxiety (e.g., fear of a burning house leads to compulsively checking electrical outlets). As part of a prefrontal circuit with the DLPFC and ACC, the RLPFC may therefore still be implicated in OCD during abstract sequencing under different task conditions.

Abstract sequential behavior differences between OCD cohorts may be due to differences in experimental conditions. A previous behavior study using this same sequential paradigm found no sequential or task switching deficits in OCD compared to HCs and persons with anxiety disorders (Doyle et al., 2024). However, there were two main experimental differences between the previous and current studies: fMRI scanning and variable (jittered for a fast event related design) inter-trial intervals (ITIs). The MRI environment, particularly noise, can cause poorer behavioral performance on cognitive tasks (Jacob et al., 2015). Further, response inhibition has been shown to be impaired in OCD compared to HCs (McLaughlin et al., 2016) and those with other anxiety disorders (Martínez-Esparza et al., 2021), which may be exacerbated by longer ITIs. It is therefore possible that lack of inhibitory control in OCD participants was more pronounced on trials with longer ITIs, resulting in increased error rates on repeat trials. However, future studies are needed to probe the effect of the MRI scanner and ITI length on behavioral performance during cognitive control tasks in OCD.

Our findings align with current neurobiological models of cognitive control in OCD. The cortico-striatal-thalamo-cortical circuits (CSTC) consist of regions shown to be dysfunctional in OCD during affective, sensory, and cognitive control tasks (Shephard et al., 2021). Specifically, the DLPFC is part of the dorsal cognitive circuit of the CSTC and is implicated in disrupting executive function, including task switching.

The ACC has been posed as a hub of information flow in the CSTC (McGovern & Sheth, 2017), with bilateral connections to the DLPFC. The DLPFC and ACC are also thought to help balance habitual and goal directed behavior in OCD (Robbins et al., 2024), with dysfunction of these regions potentially contributing to disruptions in completing goal directed tasks. Our primary results show that altered ramping dynamics in the SFS and ACC largely align with these models. Increased ramping in the SMA, MTG and TOJ informs biological models about additional cortical regions and neural dynamics implicated in cognitive and sequential control in OCD.

Potential limitations to this study are due to sample size, sample diversity, and the need for comparison to other clinical populations. Our sample contained a heterogeneous population of individuals with OCD, which limited our ability to assess symptom dimensions. The present study included individuals with comorbid anxiety diagnoses, which are common in OCD. Therefore, we were unable to directly assess the neural and behavioral effects of anxiety disorders in the present sample. The correlation between neural activity and symptom severity was relatively limited, and a larger sample size is warranted to generally investigate such potential individual differences. However, we note that despite a small sample size, we observed robust significant ramping activity in OCD compared to HCs, results which may be used to further probe the role of rACC and SFS and for future connectivity analyses to investigate contributions of networks involved in supporting abstract sequencing in OCD.

Here, we provide evidence for a neural dissociation between OCD and HCs in supporting abstract sequential and task level control. We show that increased rACC and SFS and additional cortical regions (the MTG, TOJ, and SMA) ramping supports control processes invoked during abstract sequence completion in OCD compared to HCs.

These results prompt future studies to investigate ramping dynamics in OCD and focus TMS targeting on the SFS and ACC (Grassi et al., 2023) to maintain a balance of sequential and task switching control in this population. Overall, our work highlights the importance of ramping dynamics for supporting cognitive control processes in OCD and informs neurobiological models and future treatment protocols.

## Data Availability

Imaging data are not publicly available. Analysis code is available upon request.

## Author Contributions

Hannah Doyle: Data Curation, Conceptualization, Investigation, Methodology, Formal Analysis, Writing – Original Draft, Writing – Reviewing and Editing, Visualization. Sarah Garnaat: Conceptualization, Methodology, Investigation & Training in Clinical Assessments, Clinical Resources, Writing – Reviewing and Editing, Supervision of Clinical Activities, and Funding Acquisition. Nicole McLaughlin: Conceptualization, Methodology, Investigation & Training in Clinical Assessments, Clinical Resources, Writing – Reviewing and Editing, Supervision of Clinical Activities, and Funding Acquisition. Theresa M. Desrochers: Conceptualization, Methodology, Software, Validation, Writing – Original Draft, Writing – Reviewing and Editing, Supervision, Funding Acquisition.

## Acknowledgements

We thank Dr. Ani Eloyan and members of the Desrochers Lab for their help in data collection and for their advice and feedback on the manuscript.

## References

Bergman, H., & Källmén, H. (2002). Alcohol use among Swedes and a psychometric evaluation of the alcohol use disorders identification test. *Alcohol and Alcoholism (Oxford*, Oxfordshire), 37(3), 245–251. 10.1093/alcalc/37.3.245

Berman, A. H., Bergman, H., Palmstierna, T., & Schlyter, F. (2005). Evaluation of the Drug Use Disorders Identification Test (DUDIT) in criminal justice and detoxification settings and in a Swedish population sample. European Addiction Research, 11(1), 22–31. 10.1159/000081413

Berman, A. H., Bergman, H., Palmstierna, T., & Schlyter, F. (2016). Drug Use Disorders Identification Test [Dataset]. 10.1037/t02890-000

Blanchard, T. C., Strait, C. E., & Hayden, B. Y. (2015). Ramping ensemble activity in dorsal anterior cingulate neurons during persistent commitment to a decision. Journal of Neurophysiology, 114(4), 2439–2449. 10.1152/jn.00711.2015

Bonini, F., Burle, B., Liégeois-Chauvel, C., Régis, J., Chauvel, P., & Vidal, F. (2014). Action Monitoring and Medial Frontal Cortex: Leading Role of Supplementary Motor Area. Science, 343(6173), 888–891. 10.1126/science.1247412

Bramson, B., Meijer, S., van Nuland, A., Toni, I., & Roelofs, K. (2023). Anxious individuals shift emotion control from lateral frontal pole to dorsolateral prefrontal cortex. Nature Communications, 14(1), 4880. 10.1038/s41467-023-40666-3

Chen, X., Scangos, K. W., & Stuphorn, V. (2010). Supplementary Motor Area Exerts Proactive and Reactive Control of Arm Movements. The Journal of Neuroscience, 30(44), 14657–14675. 10.1523/JNEUROSCI.2669-10.2010

Davey, J., Thompson, H. E., Hallam, G., Karapanagiotidis, T., Murphy, C., De Caso, I., Krieger-Redwood, K., Bernhardt, B. C., Smallwood, J., & Jefferies, E. (2016). Exploring the role of the posterior middle temporal gyrus in semantic cognition: Integration of anterior temporal lobe with executive processes. NeuroImage, 137, 165–177. 10.1016/j.neuroimage.2016.05.051

Demeter, Gy., Harsányi, A., Csigó, K., & Racsmány, M. (2017). STOPPING OF TASK-SWITCHING IN PATIENTS WITH OBSESSIVE-COMPULSIVE DISORDER (OCD). European Neuropsychopharmacology, 27(6), 620. 10.1016/j.euroneuro.2016.07.026

Desrochers, T. M., Ahuja, A., Maechler, M. R., Shires, J., Yusif Rodriguez, N., & Berryhill, M. E. (2022). Caught in the ACTS: Defining Abstract Cognitive Task Sequences as an Independent Process. Journal of Cognitive Neuroscience, 34(7), 1103–1113. 10.1162/jocn_a_01850

Desrochers, T. M., Chatham, C. H., & Badre, D. (2015). The necessity of rostrolateral prefrontal cortex for higher-level sequential behavior. Neuron, 87(6), 1357–1368. 10.1016/j.neuron.2015.08.026

Desrochers, T. M., Collins, A. G. E., & Badre, D. (2019). Sequential Control Underlies Robust Ramping Dynamics in the Rostrolateral Prefrontal Cortex. The Journal of Neuroscience, 39(8), 1471–1483. 10.1523/JNEUROSCI.1060-18.2018

Doyle, H., Boisseau, C. L., Garnaat, S. L., Rasmussen, S. A., & Desrochers, T. M. (2024). *Abstract task sequence initiation deficit dissociates anxiety disorders from obsessive-compulsive disorder and healthy controls*.

Etkin, A., Egner, T., & Kalisch, R. (2011). Emotional processing in anterior cingulate and medial prefrontal cortex. Trends in Cognitive Sciences, 15(2), 85–93. 10.1016/j.tics.2010.11.004

First, M. B., Skodol, A. E., Bender, D. S., & Oldham, J. M. (2017). User’s guide for the Structures Clinical Interview for the DSM-5 Alternative Model for Personality Disorders (SCID-5-AMPD) (1st ed.). American Psychiatric Publishing.

Goodman, W. K., Price, L. H., Rasmussen, S. A., Mazure, C., Delgado, P., Heninger, G. R., & Charney, D. S. (1989). The Yale-Brown Obsessive Compulsive Scale. II. Validity. Archives of General Psychiatry, 46(11), 1012–1016. 10.1001/archpsyc.1989.01810110054008

Goodman, W. K., Price, L. H., Rasmussen, S. A., Mazure, C., Fleischmann, R. L., Hill, C. L., Heninger, G. R., & Charney, D. S. (1989). The Yale-Brown Obsessive Compulsive Scale: I. Development, Use, and Reliability. Archives of General Psychiatry, 46(11), 1006–1011. 10.1001/archpsyc.1989.01810110048007

Grassi, G., Moradei, C., & Cecchelli, C. (2023). Will Transcranial Magnetic Stimulation Improve the Treatment of Obsessive–Compulsive Disorder? A Systematic Review and Meta-Analysis of Current Targets and Clinical Evidence. Life, 13(7), 1494. 10.3390/life13071494

Gu, B.-M., Park, J.-Y., Kang, D.-H., Lee, S. J., Yoo, S. Y., Jo, H. J., Choi, C.-H., Lee, J.-M., & Kwon, J. S. (2008). Neural correlates of cognitive inflexibility during task-switching in obsessive-compulsive disorder. Brain, 131(1), 155–164. 10.1093/brain/awm277

Hyafil, A., Summerfield, C., & Koechlin, E. (2009). Two Mechanisms for Task Switching in the Prefrontal Cortex. The Journal of Neuroscience, 29(16), 5135–5142. 10.1523/JNEUROSCI.2828-08.2009

Jacob, S. N., Shear, P. K., Norris, M., Smith, M., Osterhage, J., Strakowski, S. M., Cerullo, M., Fleck, D. E., Lee, J.-H., & Eliassen, J. C. (2015). Impact of fMRI Scanner Noise on Affective State and Attentional Performance. Journal of Clinical and Experimental Neuropsychology, 37(6), 563–570. 10.1080/13803395.2015.1029440

Kaldewaij, R., Koch, S. B. J., Hashemi, M. M., Zhang, W., Klumpers, F., & Roelofs, K. (2021). Anterior prefrontal brain activity during emotion control predicts resilience to post-traumatic stress-symptoms. Nature Human Behaviour, 5(8), 1055–1064. 10.1038/s41562-021-01055-2

Kauffmann, L., Ramanoël, S., & Peyrin, C. (2014). The neural bases of spatial frequency processing during scene perception. Frontiers in Integrative Neuroscience, 8, 37. 10.3389/fnint.2014.00037

Khan, A. U., Irwin, Z., Mahavadi, A., Roller, A., Goodman, A. M., Guthrie, B. L., Visscher, K., Knight, R. T., Walker, H. C., & Bentley, J. N. (2024). Low-Frequency Oscillations in Mid-rostral Dorsolateral Prefrontal Cortex Support Response Inhibition. The Journal of Neuroscience: The Official Journal of the Society for Neuroscience, 44(40), e0122242024. 10.1523/JNEUROSCI.0122-24.2024

Lapate, R. C., Ballard, I. C., Heckner, M. K., & D’Esposito, M. (2022). Emotional Context Sculpts Action Goal Representations in the Lateral Frontal Pole. Journal of Neuroscience, 42(8), 1529–1541. 10.1523/JNEUROSCI.1522-21.2021

Lashley, K. S. (1951). The problem of serial order in behavior. In Cerebral mechanisms in behavior; the Hixon Symposium (pp. 112–146). Wiley.

Lundqvist, M., Herman, P., Warden, M. R., Brincat, S. L., & Miller, E. K. (2018). Gamma and beta bursts during working memory readout suggest roles in its volitional control. Nature Communications, 9(1), 394. 10.1038/s41467-017-02791-8

Martínez-Esparza, I. C., Olivares-Olivares, P. J., Rosa-Alcázar, Á., Rosa-Alcázar, A. I., & Storch, E. A. (2021). Executive Functioning and Clinical Variables in Patients with Obsessive-Compulsive Disorder. Brain Sciences, 11(2), Article 2. 10.3390/brainsci11020267

Mayr, U., & Bryck, R. L. (2005). Sticky rules: Integration between abstract rules and specific actions. *Journal of Experimental Psychology. Learning*, Memory, and Cognition, 31(2), 337–350. 10.1037/0278-7393.31.2.337

McGovern, R. A., & Sheth, S. A. (2017). Role of the dorsal anterior cingulate cortex in obsessive-compulsive disorder: Converging evidence from cognitive neuroscience and psychiatric neurosurgery. Journal of Neurosurgery, 126(1), 132–147. 10.3171/2016.1.JNS15601

McKim, T. H., & Desrochers, T. M. (2022). Reward Value Enhances Sequence Monitoring Ramping Dynamics as Ending Rewards Approach in the Rostrolateral Prefrontal Cortex. eNeuro, 9(2), ENEURO.0003-22.2022. 10.1523/ENEURO.0003-22.2022

McLaughlin, N. C. R., Kirschner, J., Foster, H., O’Connell, C., Rasmussen, S. A., & Greenberg, B. D. (2016). Stop Signal Reaction Time Deficits in a Lifetime Obsessive-Compulsive Disorder Sample. Journal of the International Neuropsychological Society: JINS, 22(7), 785–789. 10.1017/S1355617716000540

Menon, V. (2013). Juvenile obsessive-compulsive disorder: A case report. Industrial Psychiatry Journal, 22(2), 155–156. 10.4103/0972-6748.132932

Monsell, S. (2003). Task switching. Trends in Cognitive Sciences, 7(3), 134–140. 10.1016/s1364-6613(03)00028-7

Moritz, S., Hübner, M., & Kluwe, R. (2004). Task switching and backward inhibition in obsessive-compulsive disorder. Journal of Clinical and Experimental Neuropsychology, 26(5), 677–683. 10.1080/13803390409609791

Okutucu, F. T., Kırpınar, İ., Deveci, E., & Kızıltunç, A. (2023). Cognitive Functions in Obsessive Compulsive Disorder and Its Relationship with Oxidative Metabolism. Archives of Neuropsychiatry, 60(2), 134–142. 10.29399/npa.28122

Remijnse, P. L., van den Heuvel, O. A., Nielen, M. M. A., Vriend, C., Hendriks, G.-J., Hoogendijk, W. J. G., Uylings, H. B. M., & Veltman, D. J. (2013). Cognitive inflexibility in obsessive-compulsive disorder and major depression is associated with distinct neural correlates. PloS One, 8(4), e59600. 10.1371/journal.pone.0059600

Robbins, T. W., Banca, P., & Belin, D. (2024). From compulsivity to compulsion: The neural basis of compulsive disorders. Nature Reviews. Neuroscience, 25(5), 313–333. 10.1038/s41583-024-00807-z

Rogers, R. D., & Monsell, S. (1995). Costs of a predictible switch between simple cognitive tasks. Journal of Experimental Psychology: General, 124(2), 207–231. 10.1037/0096-3445.124.2.207

Rush, A. J., Trivedi, M. H., Ibrahim, H. M., Carmody, T. J., Arnow, B., Klein, D. N., Markowitz, J. C., Ninan, P. T., Kornstein, S., Manber, R., Thase, M. E., Kocsis, J. H., & Keller, M. B. (2003). The 16-Item Quick Inventory of Depressive Symptomatology (QIDS), clinician rating (QIDS-C), and self-report (QIDS-SR): A psychometric evaluation in patients with chronic major depression. Biological Psychiatry, 54(5), 573–583. 10.1016/s0006-3223(02)01866-8

Saunders, J. B., Aasland, O. G., Babor, T. F., de la Fuente, J. R., & Grant, M. (1993). Development of the Alcohol Use Disorders Identification Test (AUDIT): WHO Collaborative Project on Early Detection of Persons with Harmful Alcohol Consumption--II. *Addiction (Abingdon*, England), 88(6), 791–804. 10.1111/j.1360-0443.1993.tb02093.x

Schneider, D. W., & Logan, G. D. (2006). Hierarchical control of cognitive processes: Switching tasks in sequences. Journal of Experimental Psychology. General, 135(4), 623–640. 10.1037/0096-3445.135.4.623

Shephard, E., Batistuzzo, M. C., Hoexter, M. Q., Stern, E. R., Zuccolo, P. F., Ogawa, C. Y., Silva, R. M., Brunoni, A. R., Costa, D. L., Doretto, V., Saraiva, L., Cappi, C., Shavitt, R. G., Simpson, H. B., van den Heuvel, O. A., & Miguel, E. C. (2021). Neurocircuit models of obsessive-compulsive disorder: Limitations and future directions for research. Brazilian Journal of Psychiatry, 44(2), 187–200. 10.1590/1516-4446-2020-1709

Tanji, J. (2001). Sequential Organization of Multiple Movements: Involvement of Cortical Motor Areas. Annual Review of Neuroscience, 24(Volume 24, 2001), 631–651. 10.1146/annurev.neuro.24.1.631

Timofeeva, P., Finisguerra, A., D’Argenio, G., García, A. M., Carreiras, M., Quiñones, I., Urgesi, C., & Amoruso, L. (2024). Switching off: Disruptive TMS reveals distinct contributions of the posterior middle temporal gyrus and angular gyrus to bilingual speech production. *Cerebral Cortex (New York*, N.Y.: 1991), *34*(5), bhae188. 10.1093/cercor/bhae188

Trach, J. E., McKim, T. H., & Desrochers, T. M. (2021). Abstract sequential task control is facilitated by practice and embedded motor sequences. *Journal of Experimental Psychology: Learning*, Memory, and Cognition, 47(10), 1638–1659. 10.1037/xlm0001004

Tsumura, K., Aoki, R., Takeda, M., Nakahara, K., & Jimura, K. (2021). Cross-Hemispheric Complementary Prefrontal Mechanisms during Task Switching under Perceptual Uncertainty. The Journal of Neuroscience, 41(10), 2197–2213. 10.1523/JNEUROSCI.2096-20.2021

Uvais, N. A., & Sreeraj, V. S. (2016). Obsessive Compulsive Disorder Presenting for Redundant Clothing. Indian Journal of Psychological Medicine, 38(1), 69–70. 10.4103/0253-7176.175126

van den Heuvel, O. A., Veltman, D. J., Groenewegen, H. J., Cath, D. C., van Balkom, A. J. L. M., van Hartskamp, J., Barkhof, F., & van Dyck, R. (2005). Frontal-striatal dysfunction during planning in obsessive-compulsive disorder. Archives of General Psychiatry, 62(3), 301–309. 10.1001/archpsyc.62.3.301

Veale, D., & Roberts, A. (2014). Obsessive-compulsive disorder. BMJ, 348, g2183. 10.1136/bmj.g2183

Zhao, Q., Wang, Z., Yang, C., Chen, H., Zhang, Y., Zeb, I., Wang, P., Wu, H., Xiao, Q., Xu, F., Bian, Y., Xiang, N., & Qiu, M. (2024). Anxiety symptoms without depression are associated with cognitive control network (CNN) dysfunction: An fNIRS study. Psychophysiology, 61(7), e14564. 10.1111/psyp.14564

